# Investigating Metabolic Trends in the Oral Cavity to Identify Novel Metabolites

**DOI:** 10.1101/2023.06.26.546600

**Authors:** Maribel E.K. Okiye, Michelle A. Velez, James Sugai, Janet Kinney, William V. Giannobile, Ashootosh Tripathi, David H Sherman

## Abstract

The human oral microbiome typically contains over 700 different microbial species. These interactions between the microorganisms within this community can shape the microenvironment throughout the human body, as these interactions are paramount to maintaining oral and overall systemic health. Recent advances in technology, such as next-generation sequencing (NGS), have revealed the complexities of the oral microbiome, linking dysbiosis of the oral microbiome with several chronic ailments such as cardiovascular disease, diabetes, and inflammatory bowel disease. However, the role of microbial secondary metabolites in oral and systemic disease progression remains poorly understood. Here, we conducted a metabolomics study on the human salivary secondary metabolome during the induction of gingival inflammation (gingivitis), the precursor to periodontal disease. In this study, we sought to assess the changes in the oral secondary metabolome during disease progression by emulating dysbiosis of the oral microbiome through a twenty-one-day induction of gingivitis in twenty human participants. We identified secondary metabolites, cyclo(L-Tyr-L-Pro) with regulatory properties for quorum sensing and inflammatory marker secretion, indicating a specialized role for secondary metabolites in oral health maintenance. Surprisingly, we also uncovered a previously unknown metabolic lag that occurs during dysbiosis recovery of the oral cavity, which indicates a lingering presence of signaling molecules for pathogenic microbe proliferation or a total oral metabolome modification following microenvironmental stress in the oral cavity. This work represents a high-resolution metabolomic landscape for understanding oral health during gingivitis that opens new opportunities for combating progressive periodontal diseases and sepsis due to the translocation of oral microbes in the human body.

---

More than a trillion microorganisms make up the human microbiome and are essential for human health and disease.^1,2^ The human microbiota, notably the gut microbiome, has been identified as an “essential organ”, playing a critical role in several biological processes such as metabolomic phenotypes, epithelial development, and innate immunity.^3–5^ However, the oral microbiome has also been linked to several systemic diseases, including obstetric convolution, cardiovascular disease, immunological disorders, diabetes, and inflammatory bowel disease.^6–8^

As the primary cross-sectional environment between the outside habitat and the internal physiology of humans, the oral cavity is considered to contain some of the most dynamic microenvironments in the body,^9^ housing nearly 800 commensal and opportunistic bacteria that constantly adapt to human lifestyle and environmental changes.^10^ The human oral microbiome is primarily inhabited by Gram-positive cocci and rods such as *Streptococcus sanguinis, Streptococcus. oralis, Streptococcus. intermedius, Streptococcus. gordonii, Peptostreptococcus micros*, and *Capnocytophaga gingivalis*.^1,11–15^ However, during the progression of diseases such as Type I and II Diabetes, there is a drastic shift in the dominant species observed, where microorganisms such as *Capnocytophaga, Porphyromonas, Pseudomonas* increase significantly in prediabetic and diabetic patients (**Table 1**).^8-12^ These changes in microbial dominance suggest nonphysical oral manifestations of systemic diseases that may contribute to or enhance disease development. The elucidation of the relationships between oral microbiota and systemic diseases is fundamentally important and will facilitate the diagnosis, prevention, and recommended treatments of the disease. Although researchers have attributed the dysbiosis of oral microbes to systemic diseases, how oral microbes mediate health and disease in the oral cavity remains unknown.

**Table 1.**
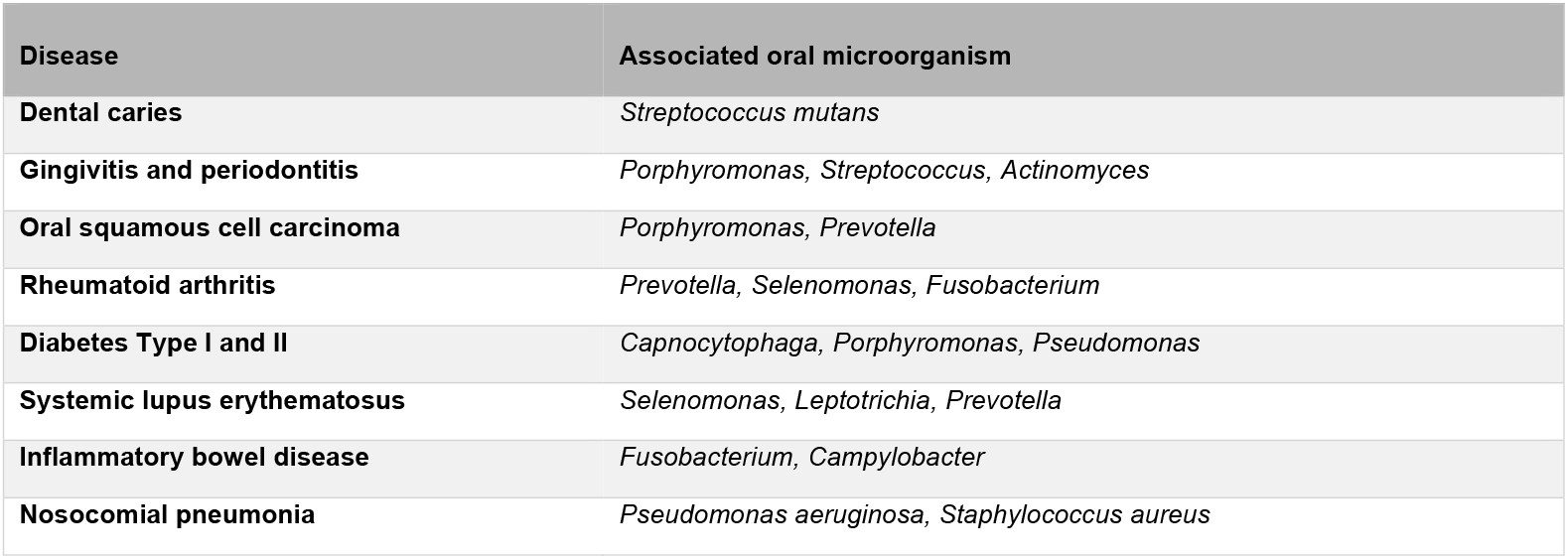
Species of oral microbes that increase in abundance during the progression of various diseases.

| Disease | Associated oral microorganism |
| --- | --- |
| Dental caries | *Streptococcus mutans* |
| Gingivitis and periodontitis | *Porphyromonas*, *Streptococcus*, *Actinomyces* |
| Oral squamous cell carcinoma | *Porphyromonas*, *Prevotella* |
| Rheumatoid arthritis | *Prevotella*, *Selenomonas*, *Fusobacterium* |
| Diabetes Type I and II | *Capnocytophaga*, *Porphyromonas*, *Pseudomonas* |
| Systemic lupus erythematosus | *Selenomonas*, *Leptotrichia*, *Prevotella* |
| Inflammatory bowel disease | *Fusobacterium*, *Campylobacter* |
| Nosocomial pneumonia | *Pseudomonas aeruginosa*, *Staphylococcus aureus* |

Saliva plays a central role in the composition, maintenance, and regulation of microbiota in the oral cavity, as it is a significant conduit for interspecies microbial communication.^16^ This is due to its complex composition as a nutrient source for supporting diverse microorganisms that maintain a healthy environment and promote plaque biofilm formation and subgingival inflammation. However, despite playing a central role in the composition, maintenance, and regulation of microbiota in the oral cavity, the salivary metabolome has not been characterized in detail compared to its corresponding proteome.^17,18^

Secondary metabolites, or natural products, are small organic compounds produced by bacteria, fungi, or plants, which are not directly involved in the organism’s normal growth, development, or reproduction.^19–22^ These specialized metabolites possess an innate affinity to cross lipid membranes and bind to diverse protein targets to exert biological functions. Secondary metabolites can mediate antagonistic interactions, such as competition and predation, and play pivotal roles as mediators of health and disease.^21–23^

In this study, we aimed to understand metabolic signatures in healthy individuals with experimentally induced gingivitis. We used untargeted metabolomics to profile secondary metabolites in the whole saliva of healthy volunteers during experimentally induced gingivitis, the precursor to periodontal disease characterized by plaque biofilm accumulation in the subgingival crevices of teeth and inflammation of the gingivae.^24–30^ The study describes a comprehensive view of metabolite dynamics during common health fluctuations in the oral cavity, giving us a new baseline for interpreting biomarkers in the oral cavity specific to gingival inflammatory disease. Moreover, we observed and identified various specialized metabolites specific to gingivitis and revealed a delay in salivary metabolome recovery. These gingivitis-specific metabolites persisted after oral hygiene resumption despite microbial abundance recovery observed in prior Experimental Gingivitis Model (EGM) studies following the standard period for resuming oral health following experimentally induced gingivitis.

## Results

### Experimental Gingivitis Model: A study of gingivitis progression in healthy volunteers

To capture the highly dynamic disease progression process at high resolution, we employed the well-established “Experimental Gingivitis Model” or EGM, developed by Löe et al. in 1965 to show correlations between taxa modifications and plaque biofilm development (**Fig. 1**).^31^

**Figure 1.**
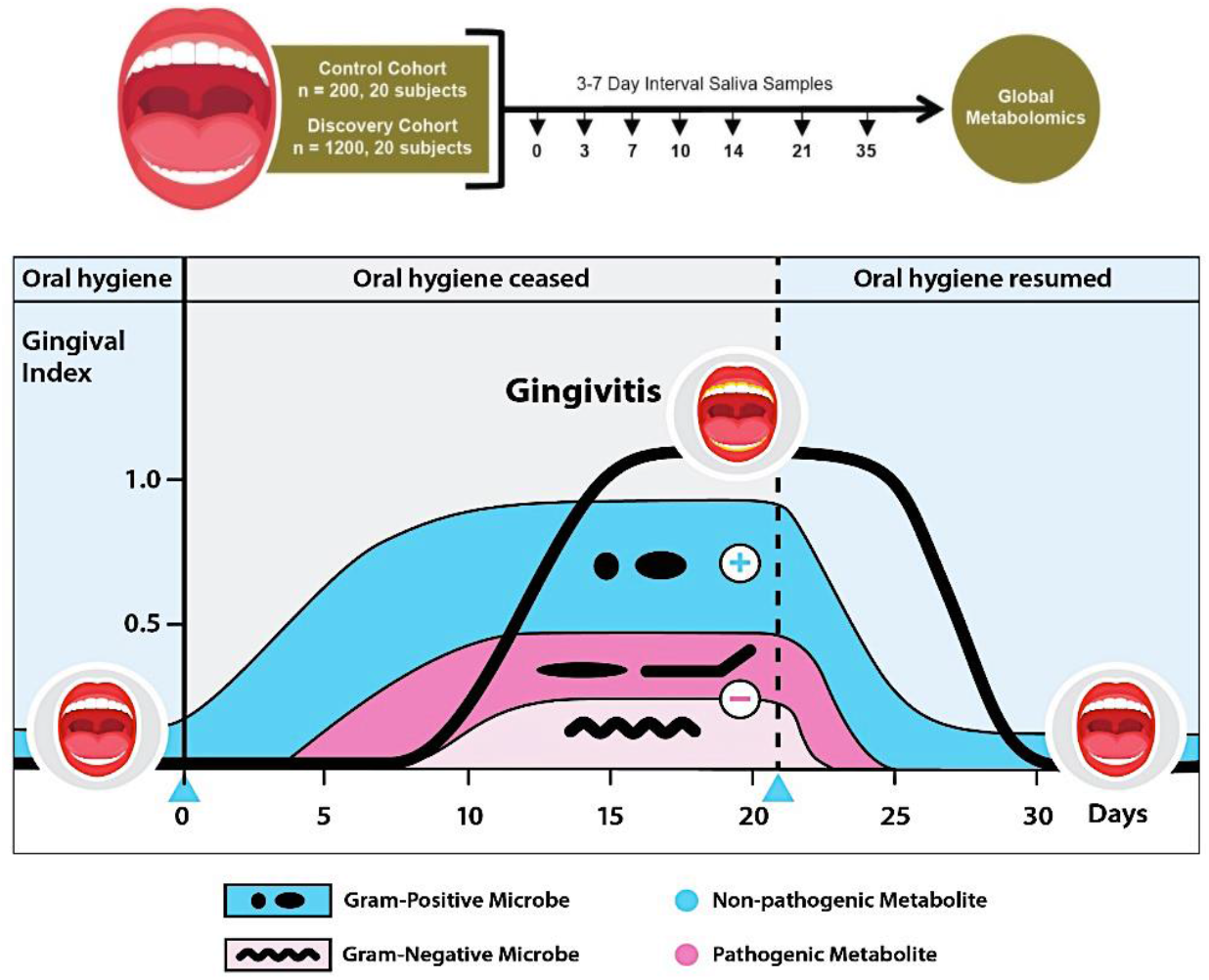
Diagram of changes in the oral cavity during the experimental gingivitis model. In a healthy oral cavity, both nonpathogenic and pathogenic microbes are in a state of symbiosis. Healthy microbes (blue) are producing metabolites for various functions and are dominant in the oral cavity. After three weeks without oral hygiene practices, plaque and gingival indices are increased, and pathogenic microbes (pink) are proliferating and producing metabolites. After the three-week mark, a professional dental prophylaxis is performed, oral hygiene practices are resumed, and the oral cavity is in the recovery phase.

Our experimental gingivitis study consisted of healthy volunteers adhering to a complete lack of oral hygiene for 21 days. This three-week period was followed by professional teeth cleaning and a 14-day recovery period (totaling 35 days, as per Löe et al. in 1965) with volunteers resuming their standard dental hygiene regimens. The timeline of our study was divided into three phases, including 1) screening phase, 2) induction phase, and 3) recovery phase **(Fig. 2)**. During this period, we measured the plaque and gingival indices and collected whole saliva samples from each subject. Participants were encouraged to continue their regular food consumption habits for more accurate visualization of oral health; however, their dietary practices and changes were not monitored. Thus, in the current study we do not account for changes in the oral cavity due to potential nutritional parameters.

**Figure 2.**
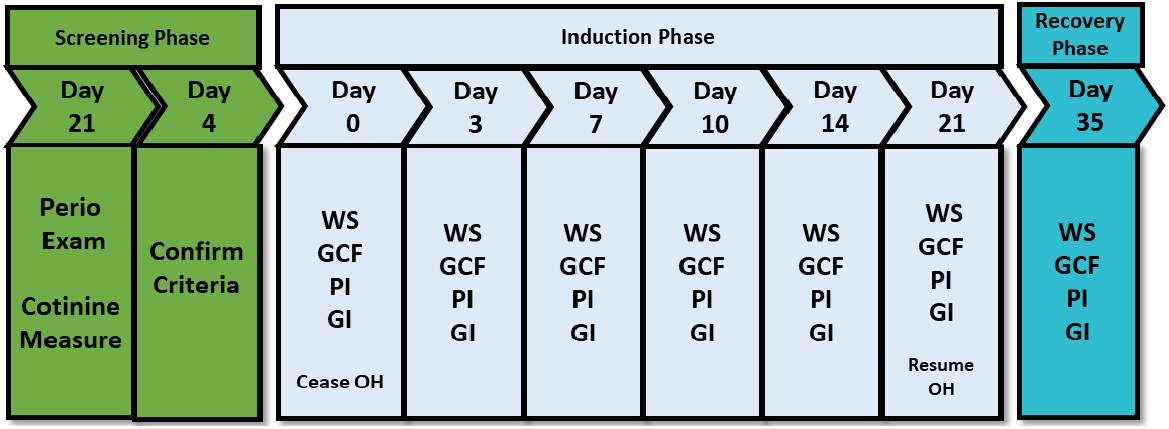
Clinical Study Timeline. Clinical study timeline with respective clinical measurements (ICF: Informed Consent Form; WS: Whole Saliva; MH: Medical History; GCF: Gingival Crevicular Fluid; PI: Plaque Index; GI: Gingival Index).

Over 300 prospective participants provided medical and dental histories were screened for potential inclusion in the investigation. An extra/intraoral comprehensive exam was performed, including a complete periodontal examination (periodontal probing depths, free gingival attachment, and clinical attachment levels). When subjects’ findings were within the study’s parameters, they were asked to provide a urine sample; and a cotinine test was performed to confirm that the participants met our “nonsmoking” specifications. Of the 300 applicants for this study only 20 met all requirements to participate. Most of our subjects were between the ages of 18 – 25 (45%), followed by 31 – 35 (30%). In addition, we had a significant number of females represented in our study (70%) but limited racial diversity, with 65% percent being Caucasian **(Table 2**).

**Table 2.** Demographic characteristics of the study volunteers.

| Characteristics |  |
| --- | --- |
| Age |  |
| 18-25 | 9 (45%) |
| 31-35 | 6 (30%) |
| 26-30 | 4 (20%) |
| 36-40 | 1 (5%) |
| Gender |  |
| Female | 14 (70%) |
| Male | 6 (30%) |
| Ethnicity/Race |  |
| Caucasian | 13 (65%) |
| Asian | 4 (20%) |
| African American | 1 (5%) |
| Bi-racial | 1 (5%) |

Whole saliva samples were collected at seven-time points throughout the 35 days on each of the 20 healthy non-smoking participants. The samples were obtained using a single-subject design with an intra-individual control (as described in Materials and Methods). Saliva samples were collected, processed, and analyzed using an untargeted metabolomics workflow.^32^ Approved volunteers for the study began EGM at t=0, where t refers to the time in days, distinguishable by the dominance of Gram-positive bacteria. At this stage, participants ceased all oral hygiene practices as noted above, including but not limited to brushing the teeth, flossing, rinsing, and chewing gum. Oral hygiene cessation was selected as the intervention for the present study because of the high reproducibility of well-established inflammatory responses in the gingival tissues. As the EGM continued, the abundance of Gram-negative bacteria increases around t=5 until t=21, when Gram-negative bacteria proliferation reaches a plateau.^33^ At t=21, the peak of induced gingivitis in the study, a professional teeth cleaning was performed, followed by the resumption of oral hygiene practices for two weeks, termed the restoration phase. Finally, at t=35, the restoration phase was complete, and the study concluded.

### Analytics of the plaque and gingival index scores

The index is defined as a numerical value describing the relative status of the population on a graduated scale with definite upper and lower limits, which are designed to permit and facilitate comparisons with other populations classified by the same criteria. In dental care, plaque and gingival indices were designed to assess oral health and patient adherence to oral hygiene protocol.^34^ The plaque index, developed by Soness and Löe in 1967 assesses plaque thickness at the crevicular margin of the teeth.^35^ Based on this thickness, dentists can provide a plaque index score” (PI score) to measure oral hygiene (**Table S1**). On the other hand, the gingival index measures the gingival status of a patient and is used to assess the severity and extent of gingival inflammation in an individual patient (**Table S1**). In tandem with metabolomic profiling, we conducted a physical analysis of each of our subject’s oral cavities by measuring the gingiva and plaque depth to assess statistically significant changes in plaque and inflammation development in oral health throughout the experimental gingivitis model period (see Table of index scores in Supplementary Table 1). These results allowed us to confirm compliance with the cessation of oral hygiene practices. Longitudinal plots were created for both GI and PI indices, along with the performance of paired-sample t-tests using IBM SPSS statistics version 26 (**Fig. S1a and S1b display the average GI and PI scores of all subjects at each time point of our study**). The entire induction and recovery phase for PI disclosed a p-value of <0.001 with a 95% confidence interval around mean PI scores for each collection day. However, the GI disclosed a p-value of < 0.001, for Day 0 to Day 21 and Day 21 to Day 35. The results for Day 0 to Day 35 did not reveal significant plaque difference (*P=*0.0764), indicating an effective removal of plaque after teeth cleaning and oral hygiene resumption. We did not use a probe to assess each collection site for the GI to avoid disrupting biofilm (e.g., plaque) formation. Despite these adaptations to PI and GI indices, statistically significant increases in PI and GI scores were observed during the induction phase, and subsequent decreases in both scores were found during the recovery phase of the study (**Fig S1a and S1b**).

### Characterization of the metabolic states of gingivitis

Previous literature regarding the experimental gingivitis model focuses on observing taxa changes in plaque as the disease progresses.^33,35–38^ However, this approach alone does not provide information on the function of individual bacteria and their role during disease progression. One way to address this limitation is by analyzing the specialized metabolites of microbes generated through diverse biosynthetic reactions.^38^ Thus, we started analyzing the alterations in oral hygiene during EGM by using untargeted metabolomics. Metabolite analysis has been regarded as the most efficient and direct approach for validating biochemical functions as it is an objective lens to view the complex nature of how physiology is linked to external events and conditions.^39^

Whole saliva metabolites were extracted using varying solvent polarities to maximize access to diverse compounds. Samples were grouped and analyzed by solvent, time-point, and subject to identify significant small molecule features using quantitative time of flight liquid chromatography-mass spectrometry (qTOF-LC-MSMS) (**Fig. S2-18**). After quality control, data filtering, and normalization, we identified over 10,059 metabolic features across the samples.^39-42^ Of these, 789 features (7.8%) were present in at least 40% of the subjects (N = 8), 56 features (9.6%) increased or diminished during gingivitis progression **(Fig. S19)**, and 26 features modulating between t=0 and t=21 with statistical significance (p>0.05 and f-change >2) (FDR = 0.017) **(Fig. 3d-e)**, suggesting consistent metabolic changes in healthy oral cavities regardless of the individual.

**Figure 3.**
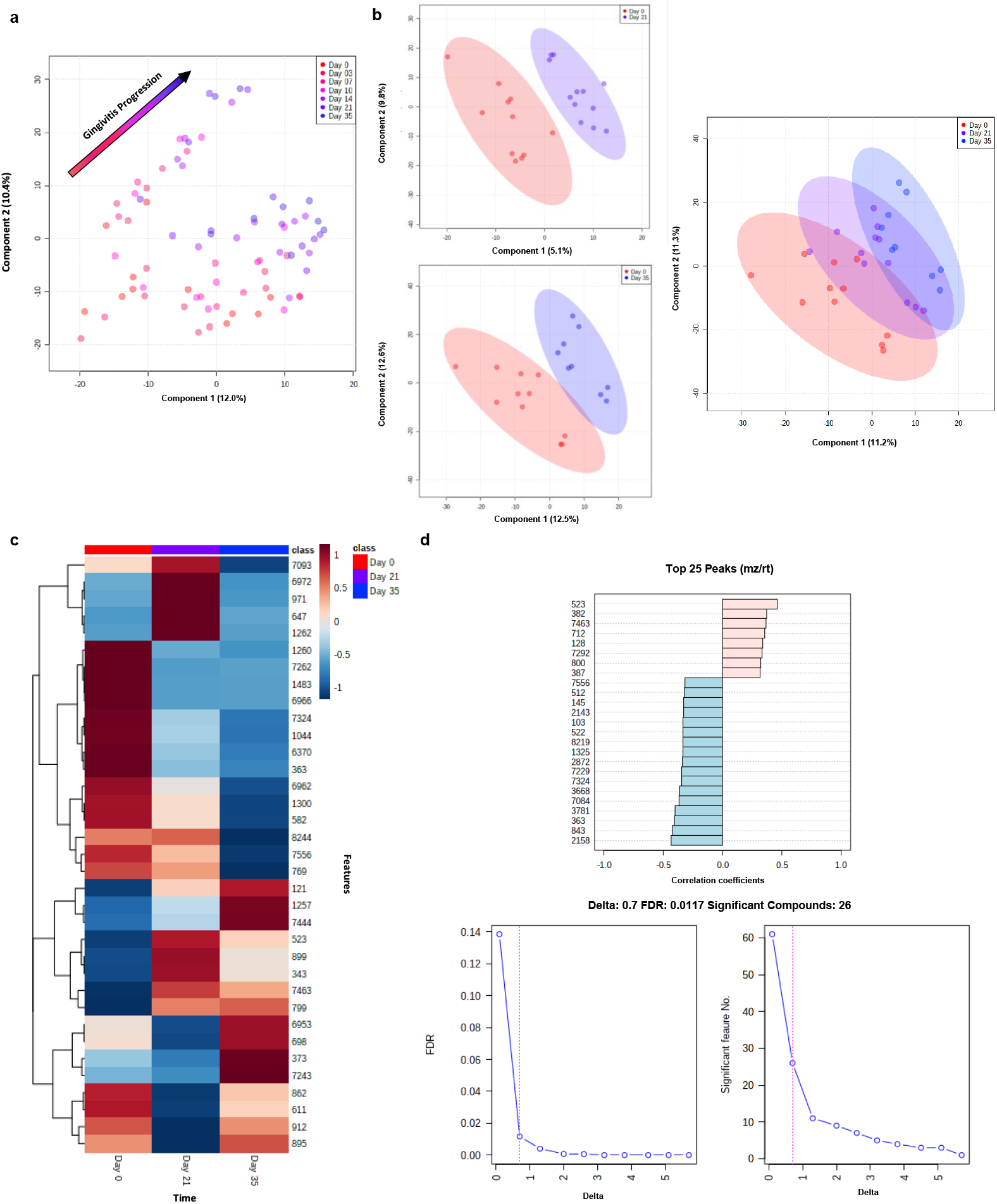
Statistical Analysis of Metabolomics Data. **a)** Partial Least Squares-Discriminative Analysis (PCA) plot model of WS from experimental gingivitis model. **b)** Partial Least Squares-Discriminative Analysis (PLS-DA) comparing t=0 vs t=21, and t=0 vs t=35 shows previously unknown variances between t=0 and t=35. The generated plot visualizes variation values and the predictive capability to discriminate between diseased and health states during induced gingivitis. **c)** Heatmap depicts the 50 most modulating metabolites between t= 0,21, and 35. **d)** The generated plot visualizes correlation coefficients between modulating metabolites and the duration of induced gingivitis in addition to the variation values and the predictive capability to discriminate between diseased and health states during induction.

We examined the metabolomic data globally using principal component analysis (PCA) and Partial Least Squares Squares-Discriminative Analysis (PLS-DA) to determine underlying metabolic trends compared to PI and GI index development. However, the global analysis of whole saliva samples via PCA could not identify metabolite trends over time (**Fig. S20**). We hypothesized this was likely attributed to variance abnormalities caused by participant-specific metabolites in the data analysis. Ultimately, we employed partial-least squares analysis to visualize molecular patterns. Metabolomics data were distributed based on the principal components 1 and 2 scores according to their stage of gingivitis regardless of individual variation and solvent extraction (**Fig. 3a)**. Interestingly, metabolites were found to have uni-directional behavior **(Fig. S21)**, with over half increasing across induced gingivitis until reaching their peak levels at t=21. While there was a gradual shift of metabolites as gingivitis progressed, with t=0 and t=3 having the most similarity, distinct metabolic patterns were observed between t=0 and t=21 (**Fig. 3b**) and t=0 and t=35 (**Fig. 3b**). We filtered our omics by subject to determine if the uni-directional behavior of metabolite modulations observed in the global analysis were conserved in inter-individual analysis. Consistent metabolite signature shifts between t=0 vs. t=21 and t=0 vs. t=35 were found across each subject (**Fig. S22a-c**). Additionally, each subject was found to have highly conserved metabolite signatures between t= 0, 3, 7, and 10, while metabolite signatures of t = 14, 21, and 35 were highly conserved. Further analysis revealed that most metabolite signature shifts occur between t=10 and t=14.

### Characterization of the oral microbiome during gingivitis progression

After analyzing the metabolic profiles of our saliva samples, we sought to connect microbial abundant shifts to the dynamic metabolic changes in human saliva using 16S RNA sequencing. A total of 35 saliva samples (seven samples from five subjects; 25% sample size) were processed for total DNA extraction (see Methods for details). We targeted, amplified, and sequenced the V3-V4 regions (approximately 460 bases) of the individual ribosomal 16S rRNA genes. An in-house developed bioinformatic pipeline was employed to process the metagenomic 16S rRNA gene sequences and generate amplicon sequence variants (ASVs) data using DADA2, QIIME 2.0, and RStudio (see Methods for details).^40^

We obtained a total of 694,859 reads that resulted in 5,055 unique ASVs using the DADA2 pipeline and strict parameters (99% similarity cutoff using SILVA SSU 138.1), with 4,636 ASVs for bacteria, and 11 ASVs for archaea at the domain level (see Methods for details including rarefaction curves and sequence quality). A total of 299 ASVs were unassigned. After filtering, denoising, and merging, the resulting ASVs were, on average 433.97 base pairs long, with 98% of sequences consisting of 466 bp total. In the bacterial domain, a total of seven established and one candidate phyla taxa were detected in 1% or more of the 35 samples, and 11 phyletic groups were found in < 1%. Approximately 3.1% of bacterial ASVs could not be further classified, indicating that a considerable proportion of novel microorganisms were detected differently from those included in bacterial reference databases. Sequencing of salivary communities across the five subjects found a total of 533 OTUs with only 125 of these present in all subjects (**Fig. S23**). This variability may partly be the result of incomplete richness coverage, moreover, comparing communities by their structure and time was more effective than comparisons based solely on membership.

The total taxonomic composition of the bacterial domain (1% cutoff), regardless of time point, depicts that the phylum with the highest number of ASVs is *Bacteroidota* (23.96% of the bacterial taxa identified in the 35 samples. Without the 1% cutoff, the *Firmicutes* phylum has the highest number of ASVs (30.83% of the bacterial taxa identified in the 35 samples). This agrees with previous studies describing the microbial composition of the oral microbiome.^31,33,36,37,41,42^ In general, Firmicutes is well distributed among subject samples, with a slightly higher abundance in subject 10 (36.22 %) compared to subject 3 (12.71 %), subject 12 (11.49 %), subject 16 (11.45 %) and subject 9 (7.44 %), (**Fig. 4a**). The next most numerous phyla are Bacteroidota (15.5 %, with Actinobacteria and Thermoleophilia as the most abundant classes), Proteobacteria (11.8 %, most of which belong to Bacteroidia), Acidobacteriota (8.6 %, with dominant levels of Vicinamibacteria and Acidobacteriae) and Chloroflexi (7.2 %, with Chloroflexia as the most prevalent class group). The 102 least frequent phyla include Cyanobacteria (3.7 %, Cyanobacteria), Myxococcota (2.3 %, Polyangia), Desulfobacterota (1.3 %, Desulfuromonadia), and candidate Methylomirabilota (1.0 %, Methylomirabilia). When comparing oral microbiomes across subjects, genera composition does not appear to vary significantly by subject. However, phylum and genera abundance does vary.

Evaluation of phylum distribution after suspension of oral hygiene revealed increases in abundance with Bacteroidata (14.4%), Firmicutes (24.2%), Fusobacteriota (5.1%), and Patescibacteria (3.8%) at the commencement of the induction phase (t=0), Bacteroidata (18.9%), Firmicutes (27.2%), Fusobacteriota (10.5%), and Patescibacteria (6.3%) and at the conclusion of induction (t=21). Then during restoration, the microbial profile returns to baseline with Bacteroidata (16.8%), Firmicutes (21.0%), Fusobacteriota (6.1%), and Patescibacteria (4.9%). Other lower-abundance phylum showing similar increasing changes in abundance included Campilobacterota and Spirochaetota. Primarily decreasing was phylum Actinobacteriota, from 13.7% (t=0), to 8.0% (t=21), and 11.7% (t=35). This pattern parallels bacterial composition shifts in dental plaque biofilm during gingival inflammation and resolution reported previously.^42^

The genus-level dynamics were driven by increases in several genus groups (Fig 5A). Although quantifications of taxonomic variations over the induction phase did not demonstrate statistically significant abundance trends, many taxonomic groups showed increasing and decreasing modulation. The most prevalent genera during induction include *Prevotella, Fusobcaterium, Veillonella*, and *Fusobacterium*, while *Streptococcus, Neisseria, Haemophilus*, and *Aggregatibacter* displayed the highest-ranking upward and downward modulating trends, consistent with the literature (**Fig 4b**). ^42^

**Figure 5.**
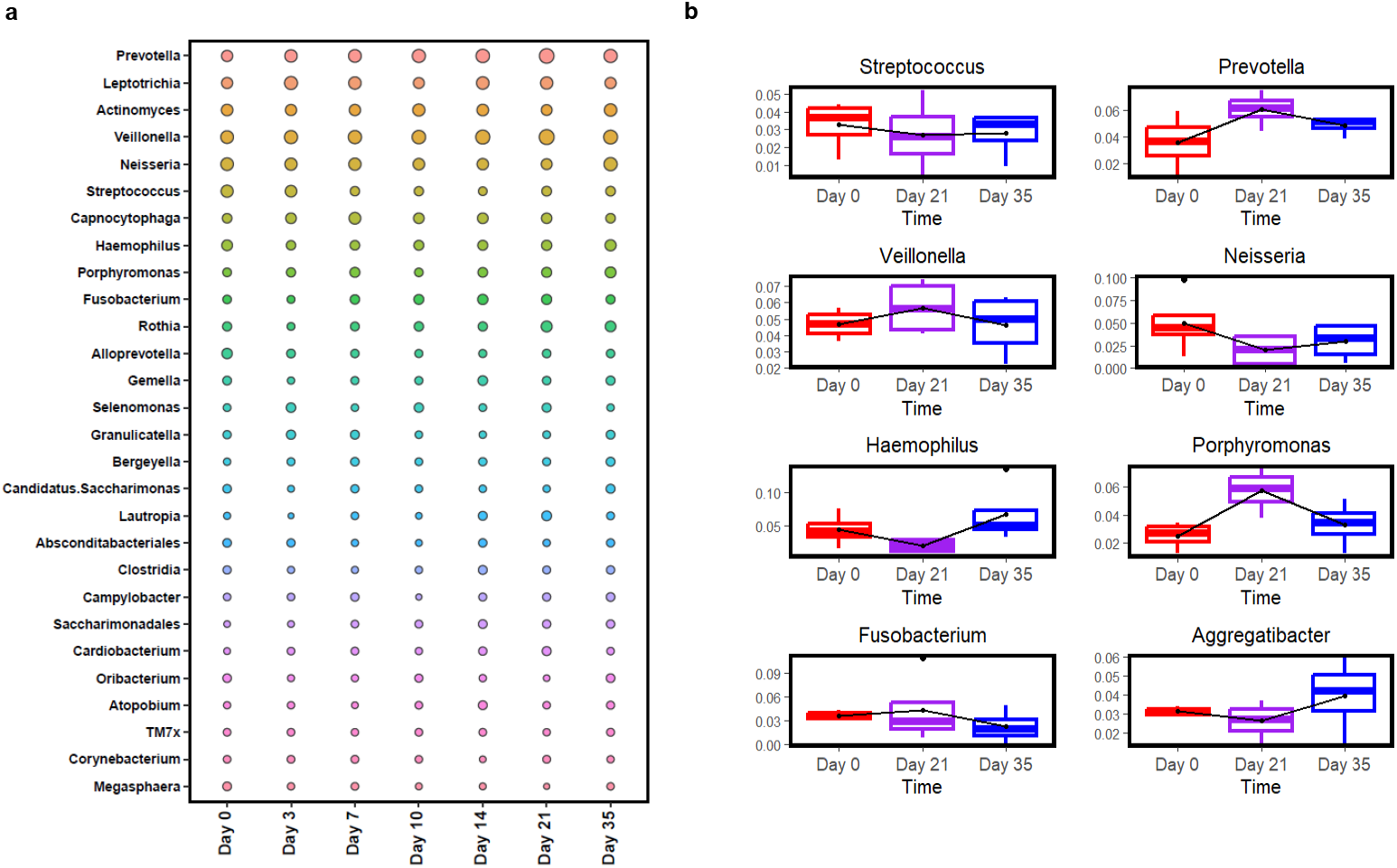
Genera distribution across samples 16S rRNA gene sequencing. Relative abundance of genera > 1% abundance in human saliva across five subjects.

**Figure 6.**
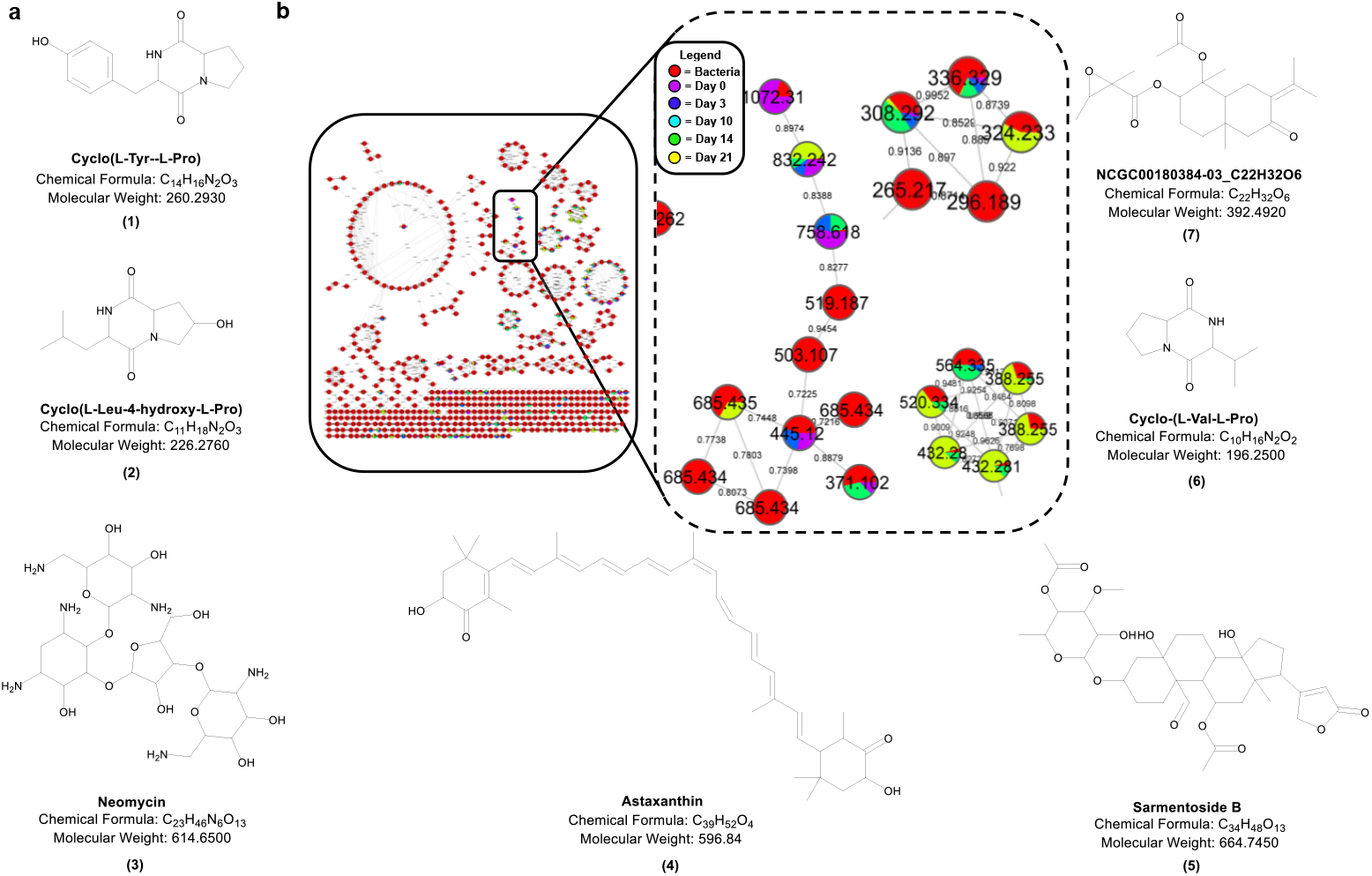
Seven annotated secondary metabolites found in human saliva correlated with oral microbe metabolites. **a)** Metabolites **(1)** cyclo(L-Tyr-L-Pro), **(2)** cyclo(L-Leu-4-hydroxy-L-Pro), **(3)** Neomycin, **(4)** Astaxanthin, **(5)** Samertoside B, **(6)** cyclo(L-Val-L-Pro), and **(7)** NCGC00180384-03 were annotated via GNPS were found in both saliva samples at various states of disease and in vitro metabolic by-products from *S. salivarius, S. mitis, S. sanguinis, and H. parainfluenzae*. **b)** The molecular network depicts 30 metabolites with high spectra similarity scores (edge_score >0.78), along with their source sample. Red indicates metabolite abundance in spectra from bacterial extracts. Yellow indicates metabolite abundance in spectra from t=0 WS samples. Green indicates metabolite abundance in spectra from t= 7 WS samples. Blue indicates metabolite abundance in spectra from t= 14 WS samples. Purple indicates metabolite abundance in spectra from t= 21 WS samples.

### Correlation of metabolomic shifts in saliva with in vitro study-based metabolite library

We then sought to identify and isolate microbially derived salivary metabolites that may have novel functions in the oral microbiome. We combined metabolomics data from the human saliva samples, and single/co-cultured oral microbe extracts to correlate metabolites modulating during gingivitis. For our *in vitro* studies, we cultured the highest-ranking oral commensals and pathogens (**Table 3**).^41,43^ Through this approach, we sought to create an extensive oral microbe metabolite library to query for secondary metabolites important in maintaining oral health. We correlated ∼320 metabolites from the human oral metabolome and conducted level 2 annotation of metabolic features by using an in-house library and public MS2 spectral databases (see details in Methods). Some of the prominent secondary metabolites identified in the samples include Sarmentoside B^44^, Diketopiperazine (DKP) cyclo(L-Tyr-L-Pro)^45^, DKP cyclo(Val-L-Pro)^46^, and the antibiotic Neomycin.^47^ (**Fig. 5a**).

**Table 3.**
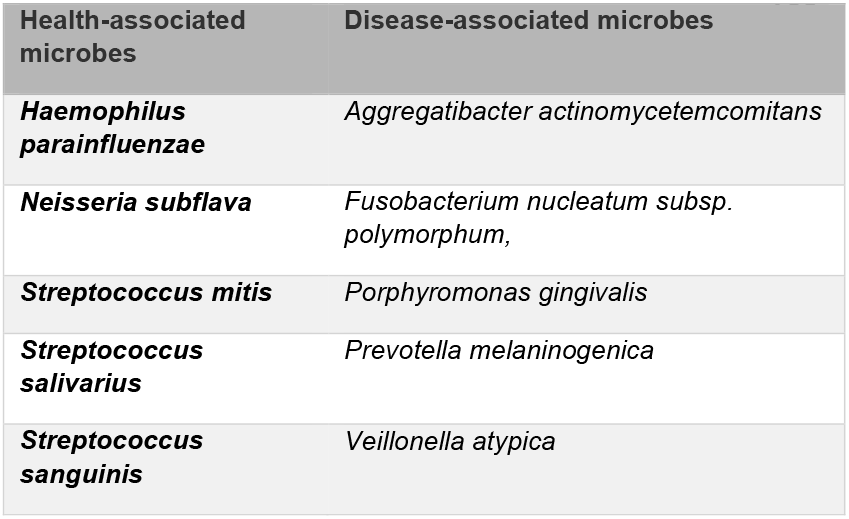
Oral microbes selected from literature studies are known to be in the healthy and diseased states of the oral cavity.

Previous work on these compounds have reported unique eukaryotic and bacterial sources and a wide variety of bioactivity. Sarmentoside B, while not originally known as a microbial metabolite, was first isolated from the plant *Strophanthus sarmentosus*.^48^ However, in 2019, it was detected in the crude extracts from a *Streptomyces cavourensis* isolated from sea cucumbers.^49^ Later, Wildenhain et al. generated a chemical-genetic matrix of 195 diverse yeast deletion strains treated with 4915 compounds and found that Sarmentoside B was predicted to have synergistic chemical-genetic interactions against at least three mutated yeast strains.^50^ Neomycin, originally isolated from the bacterium *Streptomyces fradiae*, currently is a commonly used aminoglycoside antibiotic that displays bactericidal activity against Gram-negative aerobic bacilli and some anaerobic bacilli.^51^ Annotated compounds cyclo(Val-Pro) and cyclo(Leu-Tyr) were previously evaluated under an acute renal injury model of ischemic-reperfusion.^52^ Researchers found that the two cyclodipeptides significantly abrogated ischemic injury by inhibiting renal inflammation and tubular epithelial apoptosis.^52^ Antifungal activity of cyclo(L-Pro-D-Leu) has been reported against *Aspergillus flavus* MTCC 277 and *Aspergillus niger* MTCC 282.^53^ Cyclo(L-Pro-D-Leu) was also found to strongly inhibited mycelia growth of the fungus that produces aflatoxin.^54^

DKPs are known to exhibit a variety of biological activities, most notably is their role in cell-to-cell communication used by bacteria to coordinate group behaviors.^45,46^ A mass search in our whole saliva samples revealed that DKP ions [261.1234 [*M*+H]^+^ and *197*.*8321* [*M+H]*^*+*^, which correspond to the putative diketopiperazines (DKPs) cyclo(Pro-Tyr) increased in intensity during the induction phase progression **(Fig. 5b)**. In quorum sensing, bacteria produce and respond to signaling molecules called autoinducers, which accumulate in the environment as the bacterial population grows. Once the concentration of autoinducers reaches a threshold level, the bacteria activate specific genes that control group behaviors such as biofilm formation, virulence, and antibiotic resistance. DKPs have been identified as autoinducers in several bacterial species, acting as quorum-sensing signals that regulate gene expression and promote biofilm formation.^55,56^ In addition, some studies have suggested that DKPs may interfere with quorum sensing in other bacterial species, potentially disrupting the coordination of group behaviors, and reducing bacterial virulence.^57^ With DKPs being such a promising class of natural compounds with diverse bioactivities, we sought to isolate and confirm the identities of these molecules from our oral commensals.

### Isolation and characterization of DKPs cyclo(Val-Pro) and cyclo(Tyr-Hydroxy-Pro) in oral biofilms

Based on our analysis, we sought to confirm the presence of some key metabolites using an extensive isolation and characterization methodology. We performed a large-scale fermentation of the wild-type *S. mitis* (ATCC 49456), *S. sanguinis* (ATCC BA-1455), and *S. salivarius* (ATCC 13419) at up to 10L scale. To reduce metabolite complexity, we partitioned our crude extract into fractions using reverse-phase separation followed by high-performance liquid chromatography (HPLC). The **compound (1)** was isolated as an white amorphous solid with a molecular formula of C_14_H_16_N_2_O_3_ which was derived from an HRESIMS ion peak of C_14_H_17_N_2_O_3_ [M+H]^+^ (*m/z* of 261.1251) Using the reported approximate retention time of cyclo(Tyr-Pro)(**Fig. S27**).^58^ Further analysis of ^1^H and ^13^C NMR of **(1)** suggested that it shared the same chemical shifts at position 7 suggesting the presence of tyrosine (para-hydroxyl). Further annotation confirmed the presence of at least two ketone carbons at the 1 and 10 position with chemical shifts 165.9 and 169.8, respectively, along with tertiary carbons at position 2 and 11, 59.6 and 57.2, confirming the presence of cyclo(Tyr-Pro) (**Fig S28-31**). We confirmed the annotation with previous reports as well as authentic synthetic standard (CAS #: 73-22-3) to identify of cyclo-(Tyr-Pro).^59^ (**Fig. S28-29)(Fig S30-31**).

## Discussion

This study aimed to examine the relationship between metabolome diversity and gingivitis development in the human oral cavity. Previous studies have evaluated the microbial taxa dynamics in plaque accumulation during induced gingivitis. We hypothesized that analyzing saliva meta-omics would enable a more thorough analysis of microbial changes in the oral cavity due to the complex interactions between saliva and various microenvironments. The EGM constituted a three-week gingivitis induction phase during which 20 study participants discontinued all forms of oral hygiene (brushing, flossing, mouthwash) which was followed by a final two-week recovery phase that included a full teeth cleaning, and resumption of oral hygiene practices.

Employing untargeted metabolomic profiling, we discerned a unidirectional shift of metabolites throughout the salivary metabolome, as gingivitis advanced in the oral cavity. Additionally, we determined that while distinct metabolite shifts were observed between t=0 and t=21, most metabolite signature shifts occurred between t=10 and t=14, suggesting a metabolic shift tied to competitive microbial interactions in response to the increasing abundance of pathogens associated with gingivitis. Remarkably, several metabolites associated with gingivitis, which reached their peak abundance at t=21, did not revert to their baseline levels at t=35. This suggests that the oral cavity may not parallel other microbiome systems in terms of metabolic memory, or that metabolic recovery is delayed in the oral cavity.^60^ The finding could be suggestive of epigenetic changes of the gingivae due to the microbial burden and sustained gingival inflammatory during disease induction. Due to this finding, we conducted 16S rRNA metagenomic sequencing to correlate metabolic changes to microbial compositions. While we could observe changes in the relative abundance of various oral microbes during gingivitis induction, many of these shifts were not statistically significant (P > 0.05). However, we identified trends in broad taxonomic groups **(Fig.4)** during the early onset of, and recovery from gingivitis.

The substantial modifications observed in the oral metabolite profile in this investigation provide novel insights into a high-resolution metabolomic landscape, which is fundamental in comprehending oral health alterations that occur during gingivitis. Recent studies demonstrate the continued interest in investigating panels of oral biomarkers in saliva and gingival crevicular fluid to assist in the screening and diagnosis of periodontal disease.^59^ Moreover, identifying several key metabolites associated with gingivitis progression allows researchers to distinguish early- and late-stages of gingivitis in the oral cavity. When we conducted a secondary analysis filtered by subject to assess these modulation trends in individuals, consistent metabolite signature shifts between t=0 vs t=21 and t=0 vs t=35 were found. In addition, each subject was found to have highly conserved metabolite signatures between t= 0, 3, 7 and 10, while metabolite signatures of t = 14, 21, and 35 were highly conserved. This finding reveals that ten days of induced gingivitis in the oral cavity is sufficient to change the overall metabolic profile, demonstrating that disease development initiates at the ten-day mark. These signature metabolites reveal an opportunity for metabolomic-based quantitative oral health diagnostics not previously available in dental care.

Contrary to previous literature on EGM studies, a distinct metabolite signature difference was observed between participants’ t=0 (baseline oral health) and t=35 (completion of recovery phase). The observation of a potential metabolic lag in the oral cavity, not seen at the metagenomic level, raises questions about the adequacy of current oral health practices and standards, and suggests the need for new molecular parameters. In addition, the observation contrasts with the previous EGM studies that analyzed taxa levels of plaque and found t=0 and t=35 to have similar, if not identical taxa abundance.^32^ A potential rationale for this contradiction may be related to the source of sampling analyzed. Previous evaluations of the EGM primarily focused on conducting 16S rRNA gene sequencing on plaque biomass that had accumulated on the tooth surface rather than the whole saliva. Since saliva actively moves along the mouth and contains nutrients and minerals meant to regulate the oral microbial population, saliva may be a more accurate medium to analyze changes occurring in the human oral cavity.

After successfully connecting taxa composition to metabolic by-products, we compared saliva-derived molecules to those extracted from in vitro culture, determining the sources of our microbially derived natural products. Through this, we identified 150 metabolites that were modulating during disease progression. Furthermore, we were able to connect these metabolites to a specific oral bacterium, which will provide ongoing access to metabolites of interest for future bioactivity studies. Using our analytical methods, we identified, isolated, and characterized DKP, cyclo(Tyr-Pro), that were previously reported in the literature to function as quorum sensors, indicating that our approach was successful in detecting biologically relevant compounds. We performed genome mining experiments on three bacterial strains with a known ability to produce these compounds. Unveiling the biosynthetic source of these metabolites is the subject ongoing studies.

Despite the promising findings of this study, it is important to acknowledge the relatively small sample size, which may limit the generalizability of the results. Additionally, the study was conducted in a single geographic location, which may limit the ability to extrapolate the findings to other regions, nutritional parameters or cultures. Another limitation is the lack of diversity in the study population, as the sample was predominantly composed of individuals from a single demographic group. Finally, the study relied on self-reported measures of well-being, which may be subject to bias or inaccuracies.

In conclusion, this study provides valuable insights into the relationship between metabolome diversity and gingivitis development in the human oral cavity. Our findings suggest that analyzing saliva meta-omics can offer a more comprehensive analysis of microbial changes in the oral cavity, and that distinct metabolite shifts occur during the early onset of gingivitis. These signature metabolites provide a promising opportunity for metabolomic-based oral health diagnostics not previously available in dental care. Additionally, our results reveal a potential metabolic lag in the oral cavity that raises questions about the adequacy of current oral health practices and standards and underscores the need for new molecular parameters. While this study has limitations, our approach provides a promising foundation for future research on the oral microbiome and its impact on human health.

## Acknowledgements

The protocol utilized for metagenomic data processing in this work was adapted from the metagenomic pipeline developed in the Sherman Lab (contact) in collaboration with the Universidad Nacional Agraria La Molina Lima Perú (contact)”

## Methods

### Experimental gingivitis cohort

Twenty human subjects were recruited for this study. Recruitment of subjects began after study approval was obtained. Recruitment was done through public advertising on the UM Health Research website and the posting of IRB-approved flyers at various locations on the University of Michigan Campus, including the School of Dentistry (Appendix C). All interested participants were initially screened by phone. The study recruitment approach was impartial, and all races and ethnicities were welcome to participate.

At enrollment, all participants were screened to ensure that they were healthy at baseline, without chronic conditions, and without medication intake of any kind, with no previous history of alcoholism, non-spoking, and no untreated dental issues. The medical and dental histories were updated, and blood pressure and heart rate were taken and documented. An extra/intraoral comprehensive exam was performed, including a full periodontal examination (periodontal probing depths, free gingival attachment, and clinical attachment levels). When subjects’ findings were within the study’s parameters, they were asked to provide a urine sample for a cotinine test. After a negative result, prophylaxis was performed, and oral hygiene instructions were provided. Each eligible subject was given a manual toothbrush, dental floss, and a fluoride-containing dentrifice (Colgate®) for 21 days.

### COT one-step cotinine test analysis

The COT One Step Cotinine Test Devise (Transmetron Drug Test, Salt Lake City, UT) was used to analyze the sample. A lateral flow chromatographic immunoassay was used to detect cotinine levels in human urine at a cut-off concentration of 200 mg/mL. Cotinine levels <10ng/mL are considered “negative” and consistent with no active smoking. For a subject to qualify for the study, his/her results had to be <10 ng/mL.

### Whole saliva collection and preparation

Subjects rinsed their mouths vigorously with water for 30 seconds to remove debris, then expectorated. After a two-minute waiting period to establish baseline levels of saliva, the subjects tipped their heads toward the graduated test tubes and expectorated 5mL of whole saliva in the plastic funnel inserted in the plastic, sterile tubes, labeled with the subject’s initials, sample name, and date of harvest. The sample was immediately placed on wet ice and transported to the lab. Aprotinin (1mg/ml) and Phenylmethylsulphonyl fluoride (100 mM in MeOH) were added to the whole saliva at a dilution of 1:100 and 1:200, respectively. The whole saliva was then stored at -80°C until analysis.

Within the experimental gingivitis study cohorts, 140 whole saliva samples from 20 participants were completely analyzed. Whole saliva samples were partitioned into triplicate (1.6mL per Eppendorf) and centrifuged at 10,000 rpm for 10 minutes. Pellet was separated, flash-frozen with liquid nitrogen, and saved for a further 16S rRNA metagenomic analysis. The supernatant was put through a liquid-liquid partition with three organic solvents with increasing polarity (Hexanes; Ethyl Acetate; Butanol). The four extractions (BtOH, EtOAc, Hexanes, and AQ) were dried under vacuum (Biotage V10 Touch) and resuspended in 100% MeOH (BtOH; EtOAc; AQ Layer) and MeCN 100% (Hexanes). Metabolic Extracts were then analyzed via LC-MS/MS.

### Strains and culture conditions

*Fusobacterium nucleatum* subsp. *polymorphum* (ATCC 10953), *Prevotella melaninogenica* (ATCC 700524), *Porphyromonas gingivalis* (ATCC 33277), and *Veillonella parvula* (ATCC 17745) were maintained short-term on BD Brucella Blood Agar with Hemin and Vitamin K1 (Cat. No. 255509). Starter cultures were grown in reinforced clostridial medium (Oxoid CM0149) with 1.5% v/v 60% sodium lactate, and 4mM filter sterilized sodium sulfite. Plates and liquid cultures were grown in anaerobic atmospheric conditions consisting of 85% N_2_, 10% CO_2_ and 5% H_2_. *Veillonella parvula (*ATCC 17745), kindly donated by Dr. Erik Krukonis, University of Detroit – Mercy. *Streptococcus mitis (*ATCC 49456*), Streptococcus salivarius (*ATCC 13419*), Streptococcus sanguinis (*ATCC BA-1455*), and Neisseria subflava (*ATCC 49275) were maintained on BHI agar and grown in liquid BHI and grown in aerobic atmospheric conditions with 10% CO_2_. Cultures of the previous microorganisms were incubated in an anaerobic atmosphere consisting of 5% H_2_, 10% CO_2_, and 85% N_2_.

### MS acquisition and chromatographic conditions

Metabolic extracts were analyzed by LC-HRMS/MS in positive ionization modes. Agilent 1290 Infinity II UPLC coupled to an Agilent 6545 ESI-Q-TOF-MS system was operated in auto MS/MS-scan mode for data acquisition (acquisition from m/z 100 to 2,000) with an MS scan rate of 5.00 scans/sec and MSMS scan rate of 3.00 scans/sec with the resolution set at 30,000 (at m/z 400). Chromatography was performed using a Phenomenex Kinetex 5 μm Phenyl-Hexyl 100 Å column (50 × 4.6 mm). Compound elution was accomplished utilizing a gradient starting with a 1 min isocratic step consisting of 90% A (A: 100% H2O followed by a 7 min linear gradient elution to 100% B (100% MeCN), followed by an isocratic plateau step consisting of 100% B for 2 min (flow rate of 0.4 mLmin-1). The divert valve was set to waste for the first 1 min. Samples were injected at a concentration of 100 μgmL-1 (5 μL), and elution was detected by ESI - MS, and UV 195−400 nm. All reagents were purchased from commercial sources and used without further purification. All solvents used for compound purification were HPLC grade unless otherwise stated. The resulting mass spectra were exported into Mzmine 2. MZmine (V4.5.2, obtained from the project website at: http://mzmine.sourceforge.net/.) was used to analyze the LC-MS raw data, with orthogonal partial least-squares discriminant analysis (OPLS-DA) in the MetaboAnalyst 5.0 web server (http://www.metaboanalyst.ca, accessed on 24 March 2022) and SIMCA (V.14.0, Umetrics Inc., Umea, Sweden.) software. A VIP > 1, the fold change (FC) value (FC > 2 and FC < 0.5), and a *t*-test (*p* < 0.05) were used to determine the differential metabolites.

### Quantification and Statistical Analysis

#### Section 1: Metabolomics Data Pre-processing

After conversion to .mzML using MSConvertGUI v3.0 (ProteoWizard Software Foundation, Palo Alto, CA, USA), raw files were pre-processed and features were detected using MZmine v2.53. Data were cropped based on retention time (RT) 1–9. min. Masses were detected with a noise threshold of 10,000 for MS1 and 100 for MS2. The chromatogram was built using the ADAP module, with minimum 5 scans per peak, a group intensity threshold of 15,000, a minimum highest intensity of 40,000, and a *m/z* tolerance of 0.002 *m/z* or 10 ppm. Deconvolution was performed using the ADAP module, with *m/z* center calculation using median, and ranges for MS2 scan pairing of 0.1 Da and 0.2 min. The isotopes were grouped with a *m/z* tolerance of 0.002 *m/z* or 10 ppm and RT tolerance of 0.1 min. Peaks were aligned with a *m/z* tolerance of 0.002 *m/z* or 10 ppm and RT tolerance of 0.1 min, with 75% weight given to *m/z* and 25% to RT. Finally, peaks were filtered with a minimum of 7 peaks in a row. The feature quantification table (.csv) and aggregated MS2 mass list (.mgf) were exported for further analysis using Metaboloanalyst.

#### Section 2: Metabolic Features Identification

Metabolite identification was performed using a two-step approach. First, to identify compounds, we used our in-house metabolite library from the Natural Product Discovery Core, which contains chemical standards and a manually curated compound list based on accurate mass (m/z, ± 10 ppm), retention time and spectral patterns. Second, further metabolites were identified based on accurate mass, isotope pattern and MS/MS spectra against public databases, including HMDB, MoNA, MassBank, <u>METLIN,</u> and NIST. The generated MS1/MS2 pairs were searched in the public databases: HMDB, MoNA, and MassBank. The MS/MS spectra similarity score was calculated using the forward dot-product algorithm, which considers both fragments and intensities. The similarity score cutoff was set as 0.5. The metabolic features with MS/MS spectra that were not matched in downloaded public databases were searched in the online public databases, METLIN and NIST. Then the MS/MS spectra match was manually checked to confirm the identifications (Level 2 Identification).

#### Section 3: Identification of Significantly Altered Features/Compounds

In order to conduct statistical analyses for untargeted metabolomics, MetaboAnalyst (v5.0) was utilized. Compound abundances were normalized by median, and log transformed without scaling. A partial least square-discriminant analysis (PLS-DA) was conducted to putatively characterize patterns between time points across gingivitis progression. Supervised Pearson correlation heatmaps and Ward’s hierarchal clustering were also conducted for the most significant 50 metabolic relationships based on variable importance in projections (VIP) values from PLS-DA analyses for whole saliva samples. A statistical method specialized for multi-testing, significance analysis of microarrays) was applied to identify metabolic features/compounds altered significantly in metabolome-wide analysis. Specifically, we used SAM to examine the correlation between abundance of each compound and the time into disease progression of each sample. For all SAM analyses, distribution-independent ranking tests (based on the Wilcoxon test) and the sample-wise permutation (default by the samr package) were used to ascertain significance (false discovery rate, FDR < 0.05).

