## Supporting Information for "Investigating Metabolic Trends in the Oral Cavity to Identify Novel Metabolites"

**Supplementary Information**

| Scores | Plaque Index (PI) Criteria | Gingival Index (GI) Criteria |
|---|---|---|
| 0 | No plaque | Absence of inflammation |
| 1 | A film of plaque adhering to the free gingival margin and adjacent area of the tooth. The plaque may be seen *in situ* only after the application of disclosing solution or by using the probe on the tooth surface. | Mild inflammation A slight change in color and a little change in texture. |
| 2 | Moderate accumulation of soft deposits within the gingival pocket, or on the Tooth and gingival margin, which can be seen with the naked eye. | Moderate inflammation Moderate glazing, redness, edema, and hypertrophy. Bleeding on pressure. |
| 3 | The abundance of soft matter within the gingival pocket and/or on the tooth and gingival margin. | Severe inflammation Marked redness and hypertrophy. Tendency to spontaneous bleed. Ulceration. |

**Supplementary Table 1.** Indices used for clinical investigation.

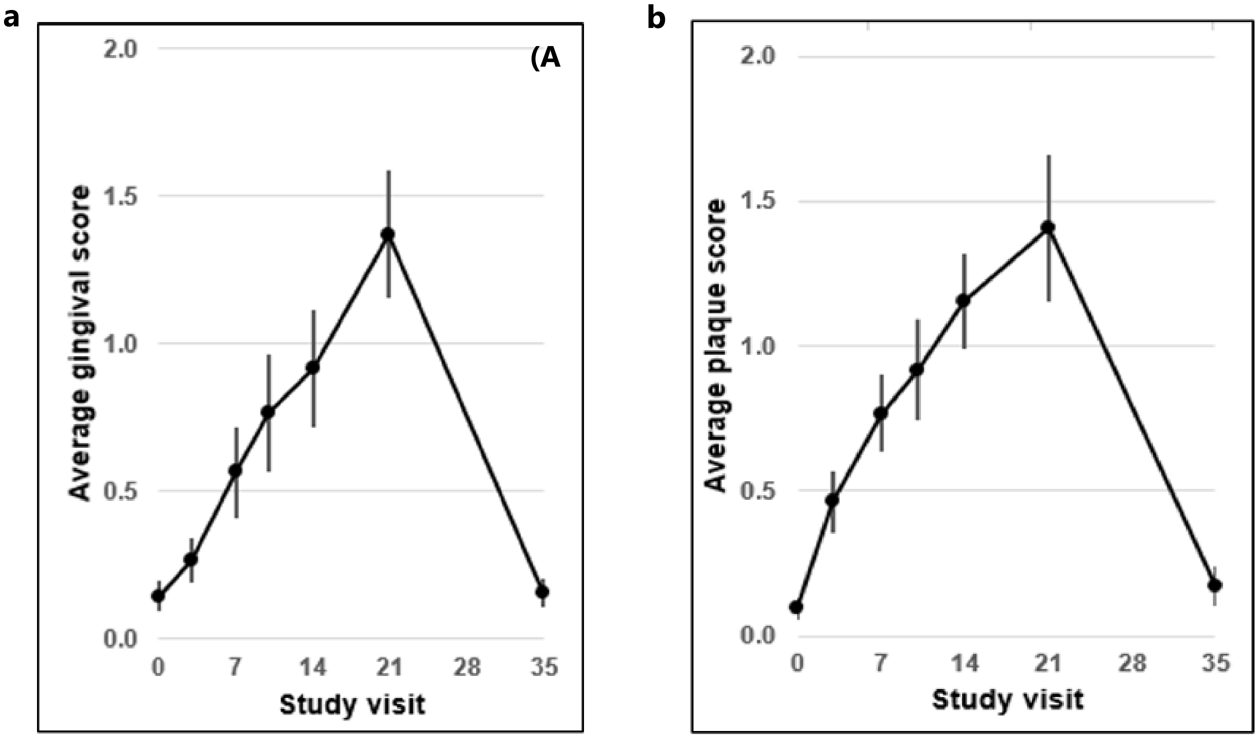


**Supplementary Figure 1. Physical Indications of Gingivitis.** Longitudinal PI and GI represent study participants' pre-induction, induction, and recovery phase scores (Day 3-Day 35). Progression. Diagram depicts the increasing (Day 0 – 14) and decreasing (Day 21-15) clinical means for **(a)** gingival and **(b)** plaque indices of the subjects throughout induced gingivitis.

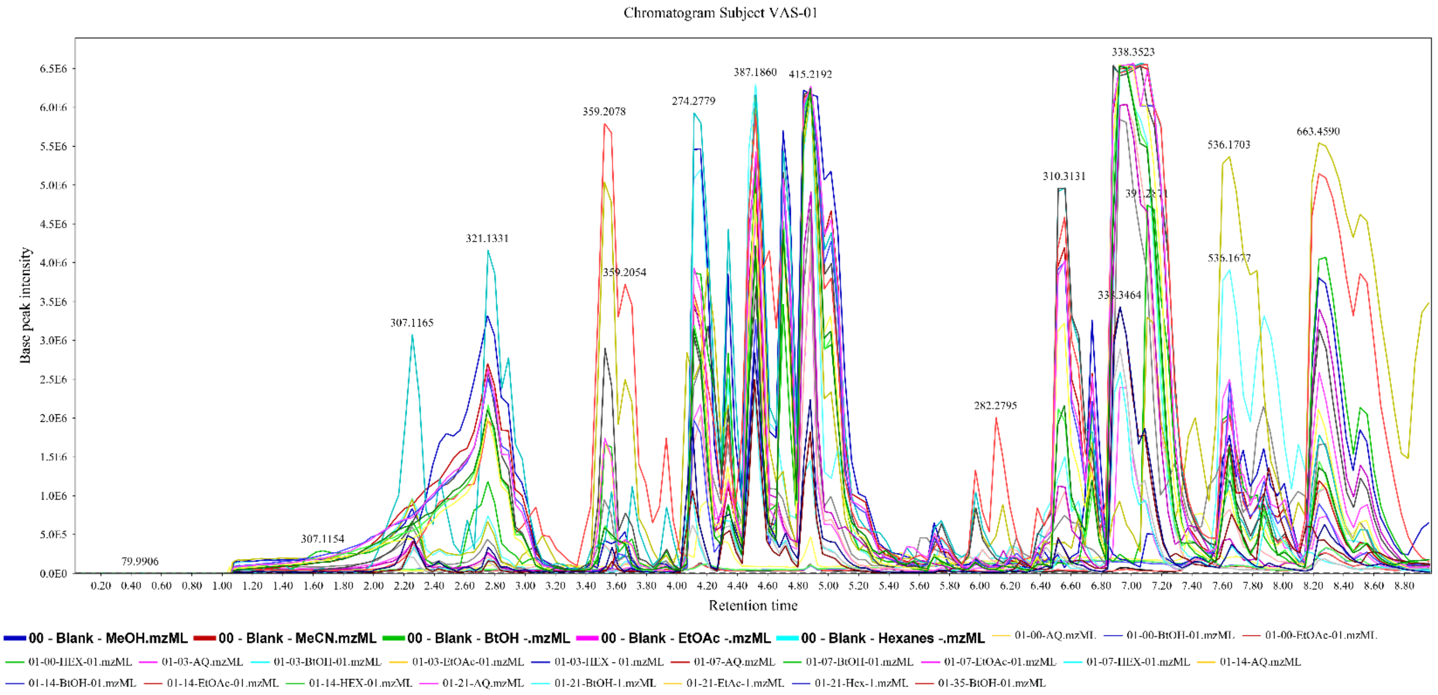


**Supplementary Figure 2. HPLC Chromatogram of Whole Saliva from Subject VAS-01 during Induction and Recovery.**

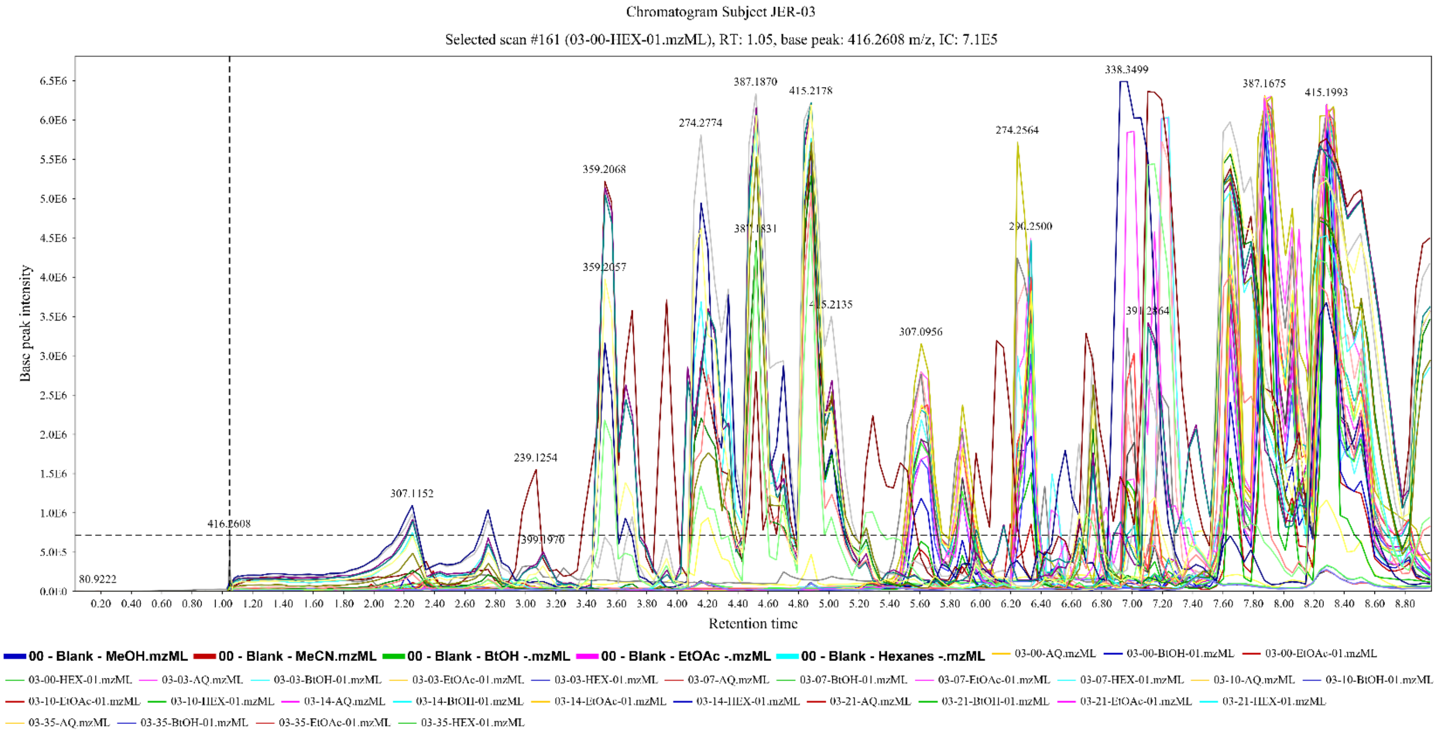


**Supplementary Figure 4. HPLC Chromatogram of Whole Saliva from Subject JER-03 during Induction and Recovery.**

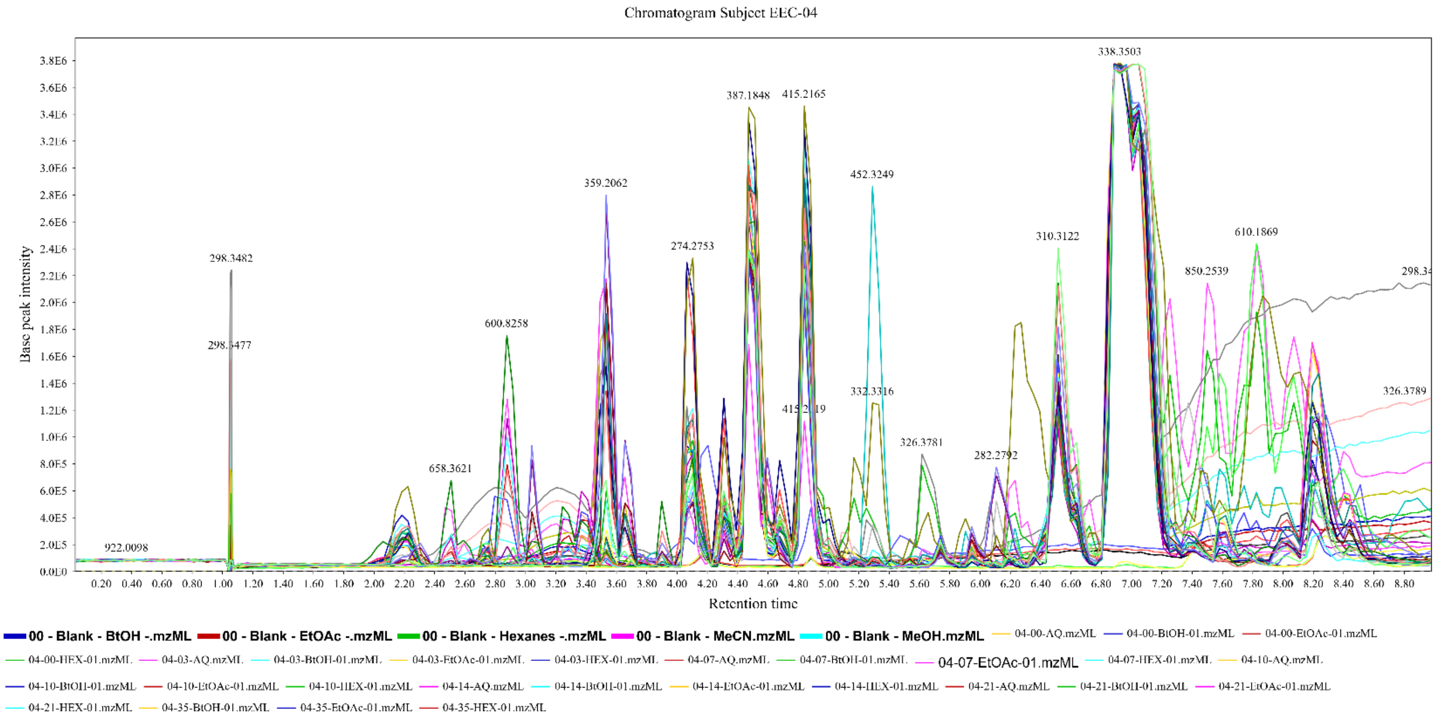


**Supplementary Figure 5. HPLC Chromatogram of Whole Saliva from Subject EEC-04 during Induction and Recovery.**

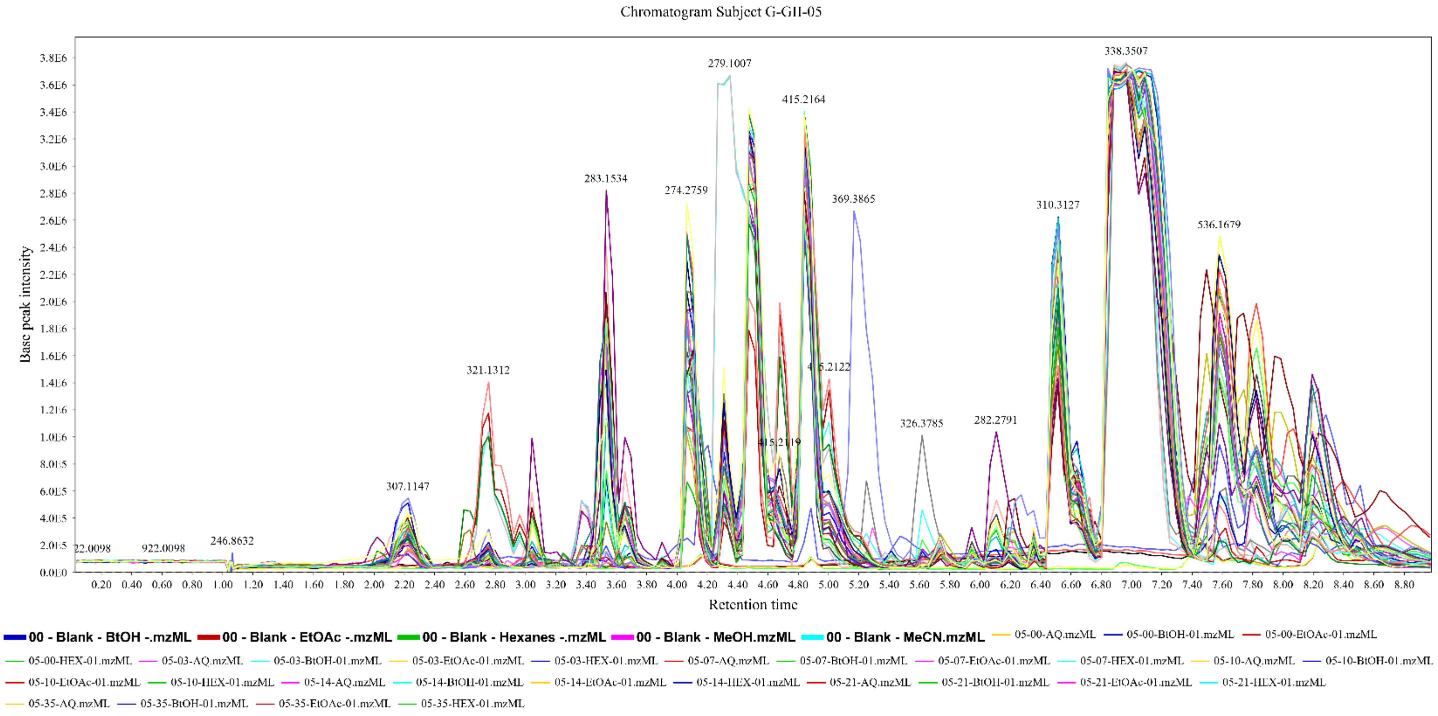


**Supplementary Figure 6. HPLC Chromatogram of Whole Saliva from Subject G-GII-05 during Induction and Recovery.**

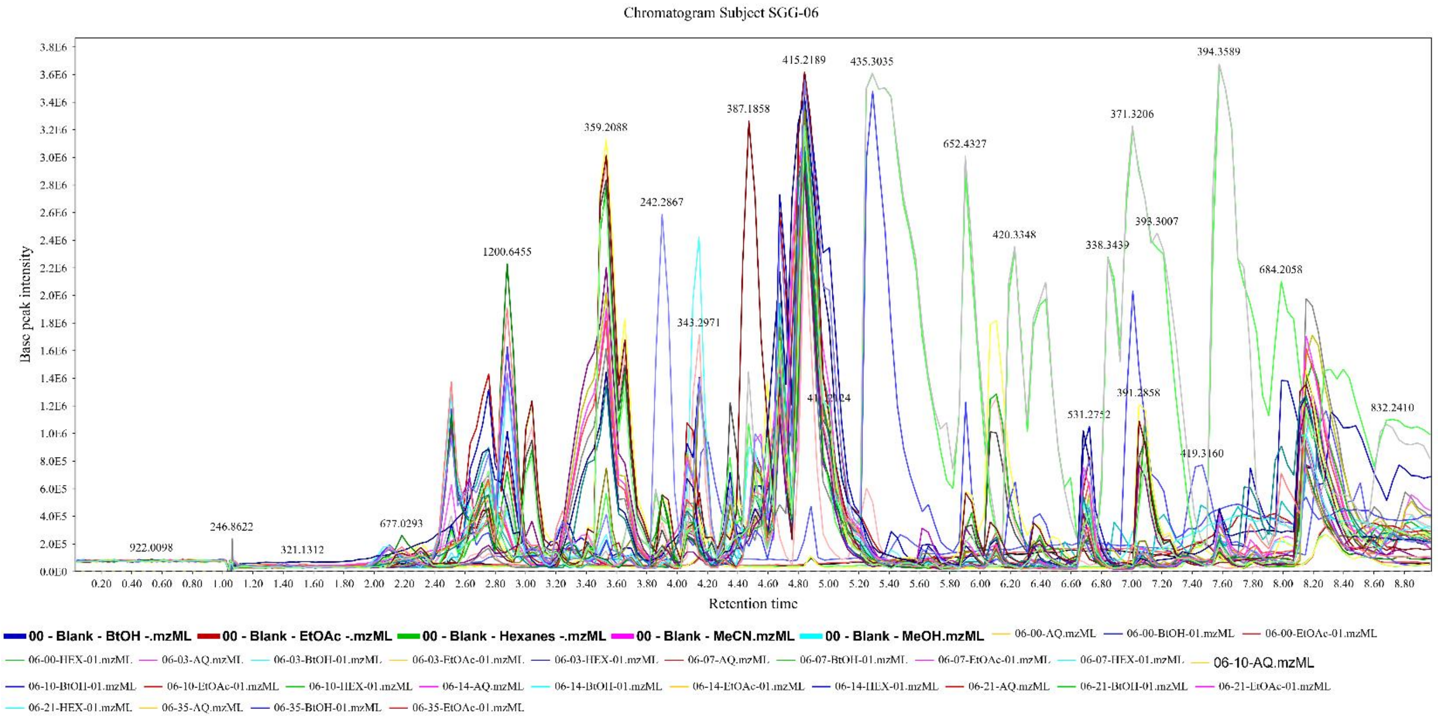


**Supplementary Figure 7. HPLC Chromatogram of Whole Saliva from Subject SGG-06 during Induction and Recovery.**

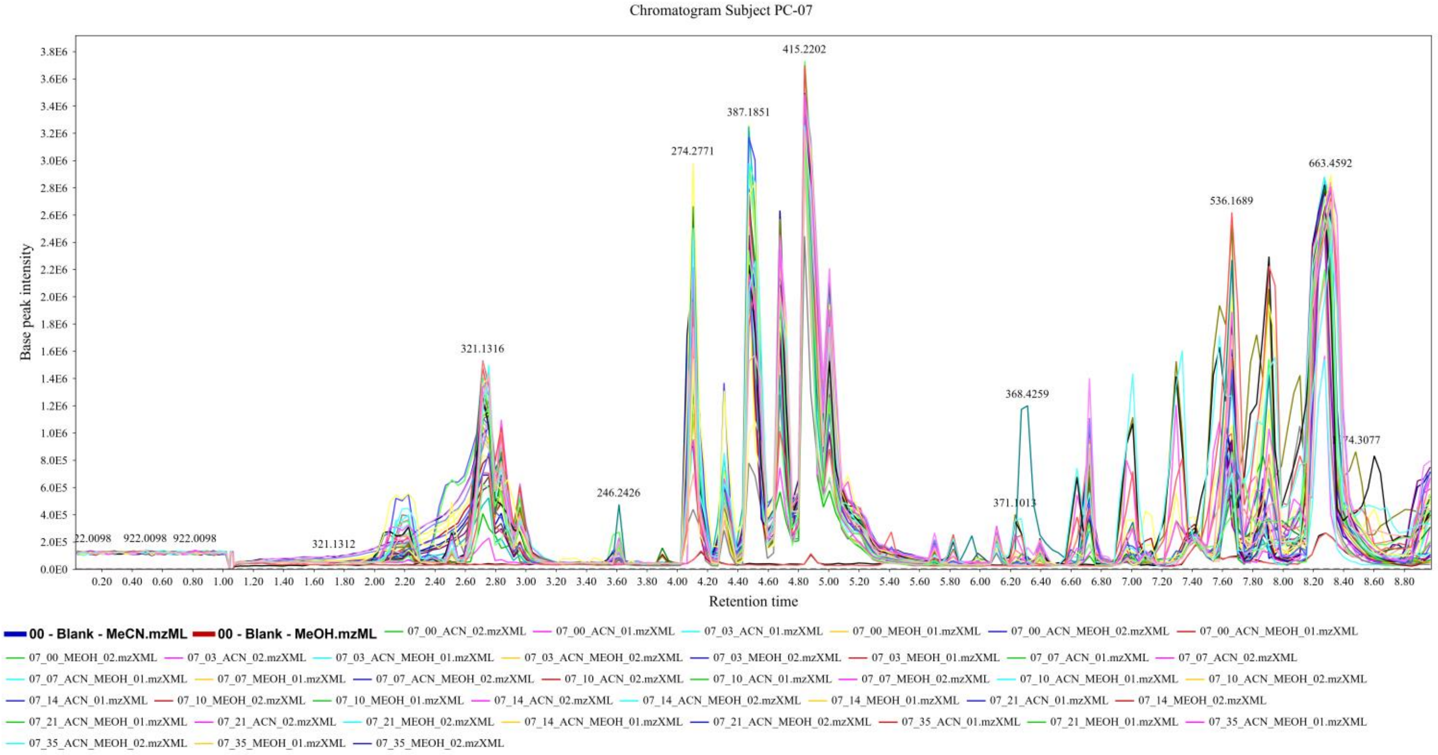


**Supplementary Figure 8. HPLC Chromatogram of Whole Saliva from Subject PC-07 during Induction and Recovery.**

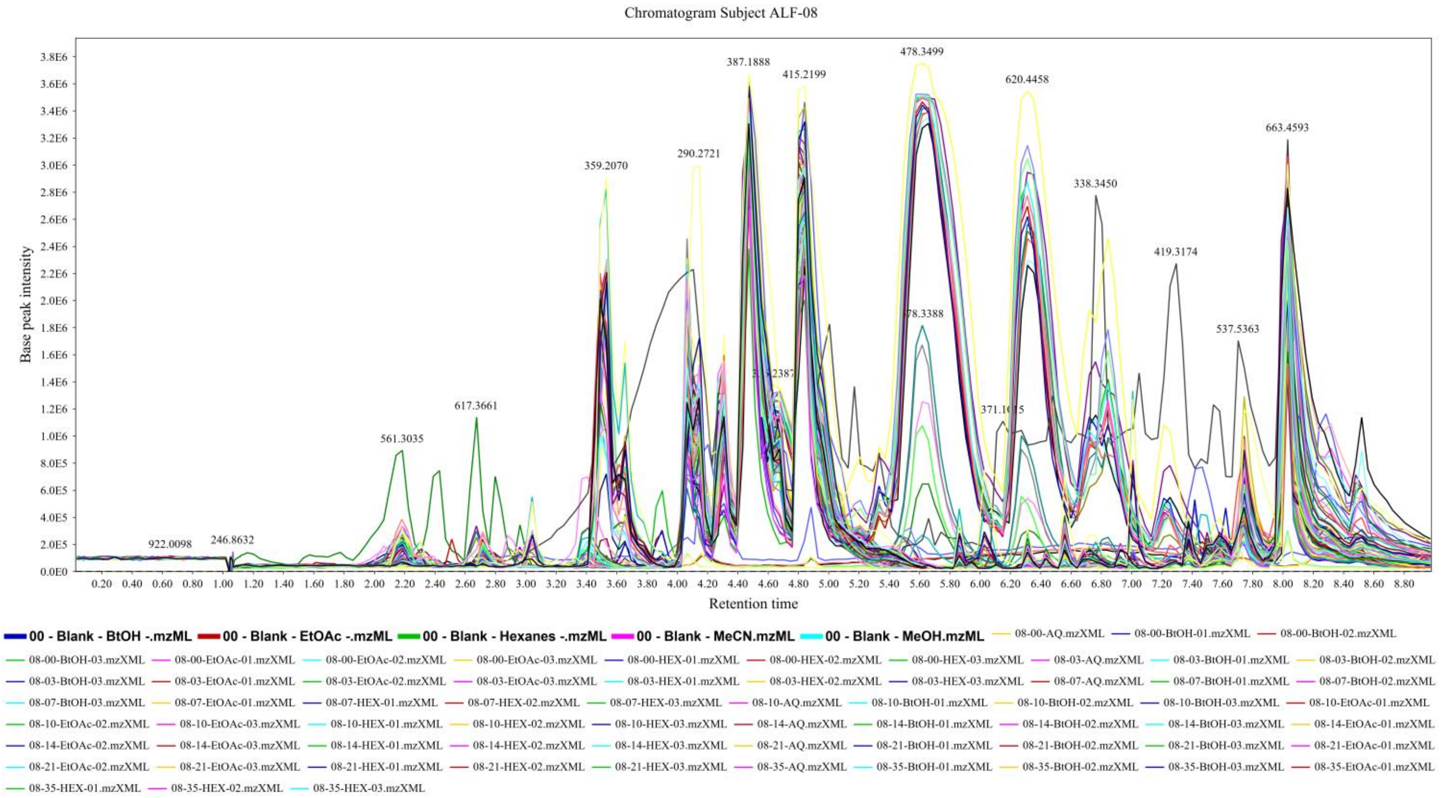


**Supplementary Figure 9. HPLC Chromatogram of Whole Saliva from Subject ALF-08 during Induction and Recovery.**

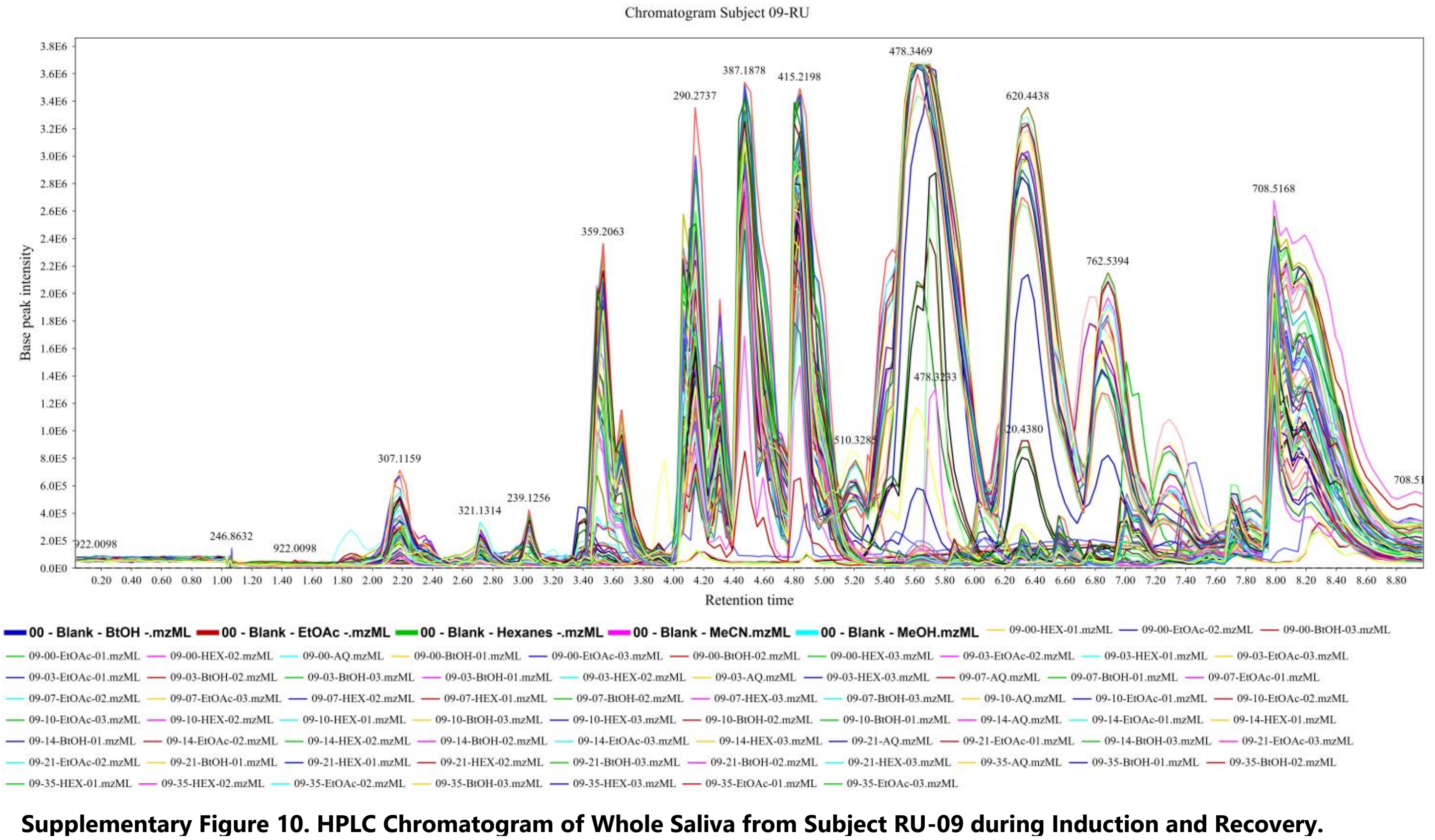


**Supplementary Figure 10. HPLC Chromatogram of Whole Saliva from Subject RU-09 during Induction and Recovery.**

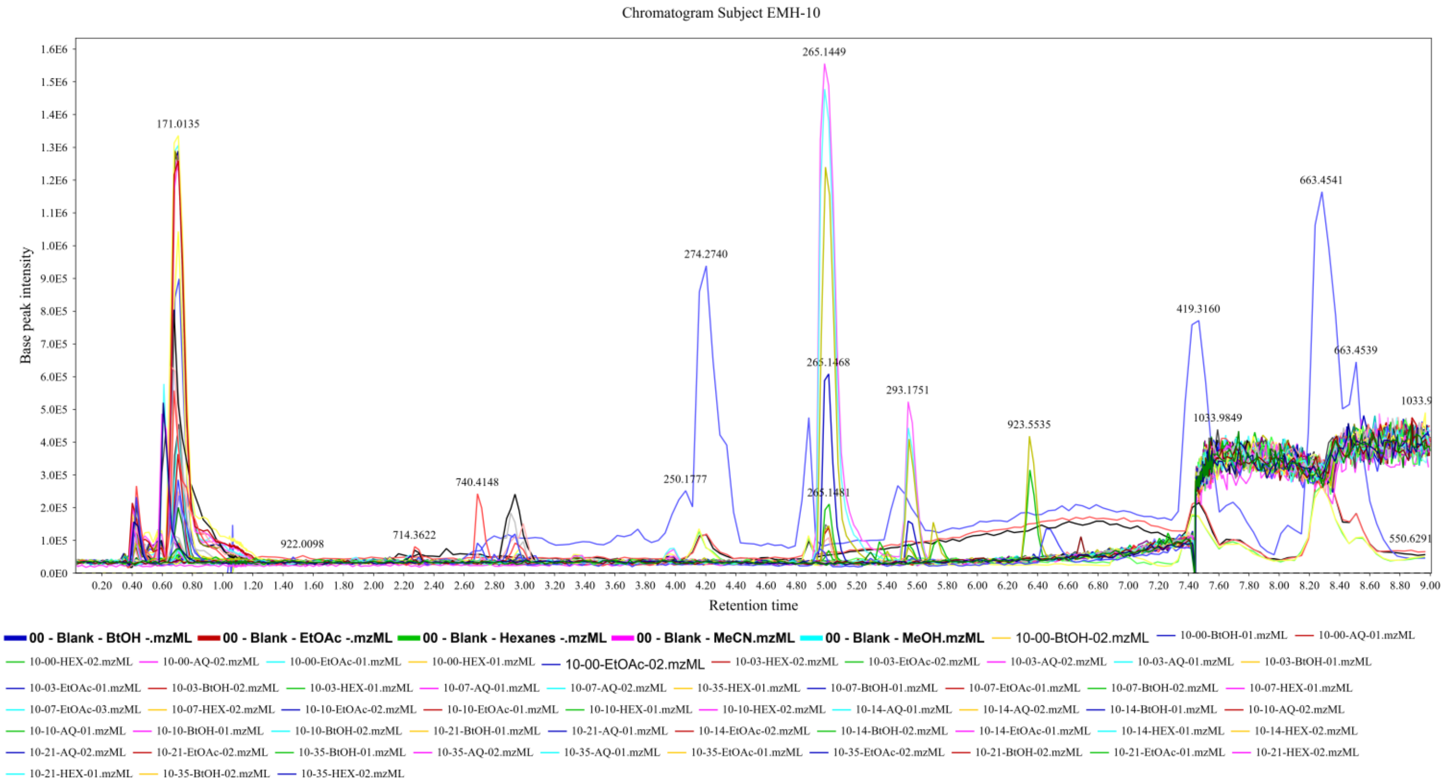


**Supplementary Figure 11. HPLC Chromatogram of Whole Saliva from Subject EMH-10 during Induction and Recovery.**

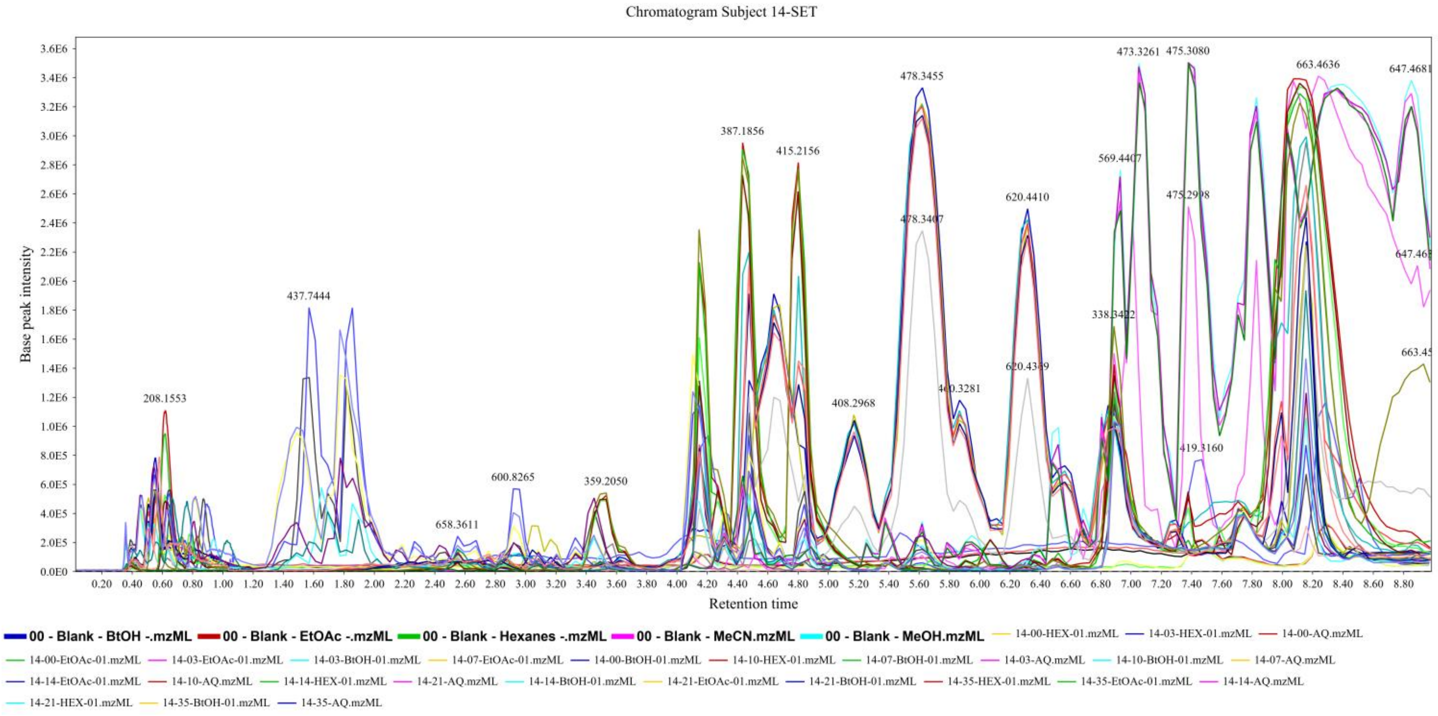


**Supplementary Figure 12. Supplementary Figure 10. HPLC Chromatogram of Whole Saliva from Subject -11 during Induction and Recovery.**

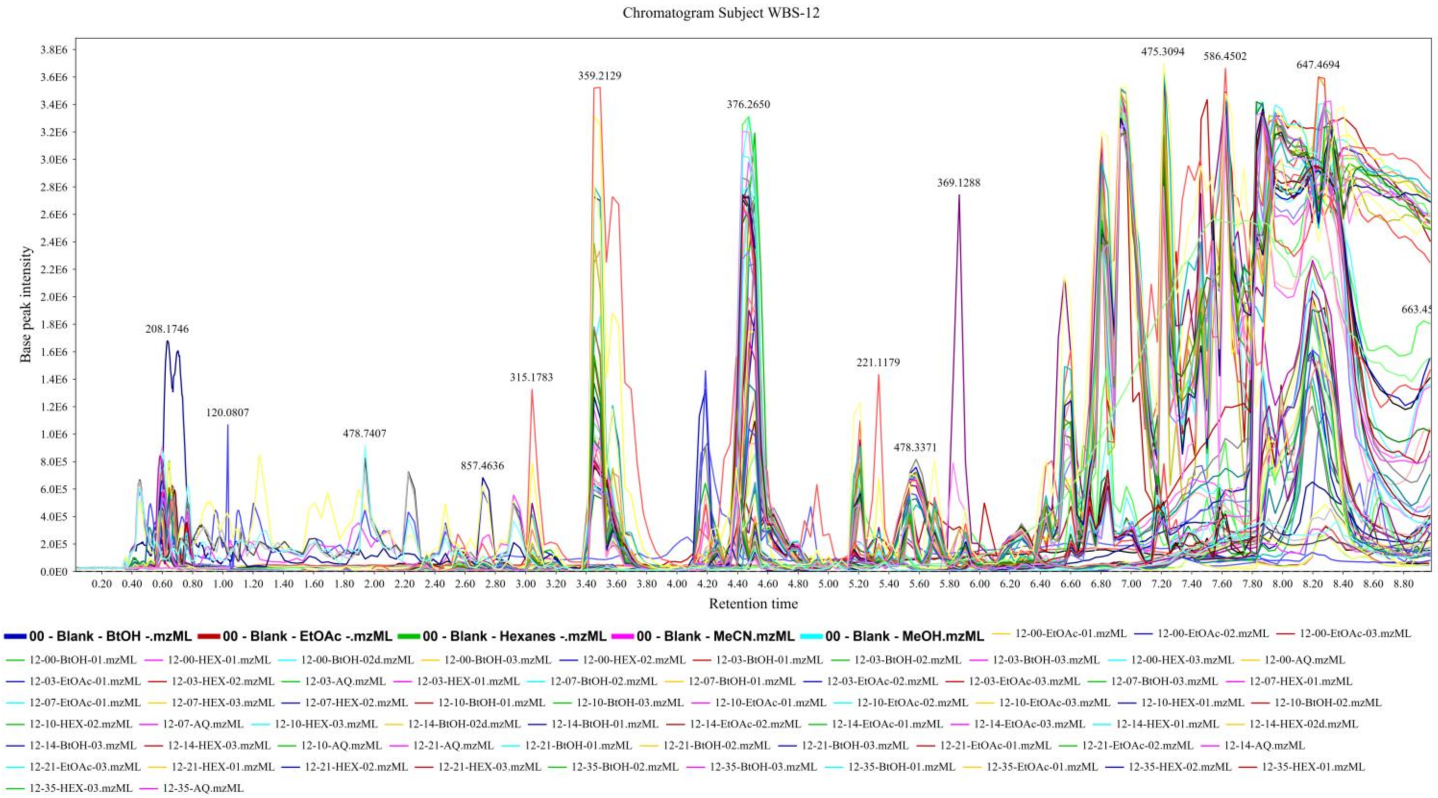


**Supplementary Figure 13. HPLC Chromatogram of Whole Saliva from Subject WBS-12 during Induction and Recovery.**

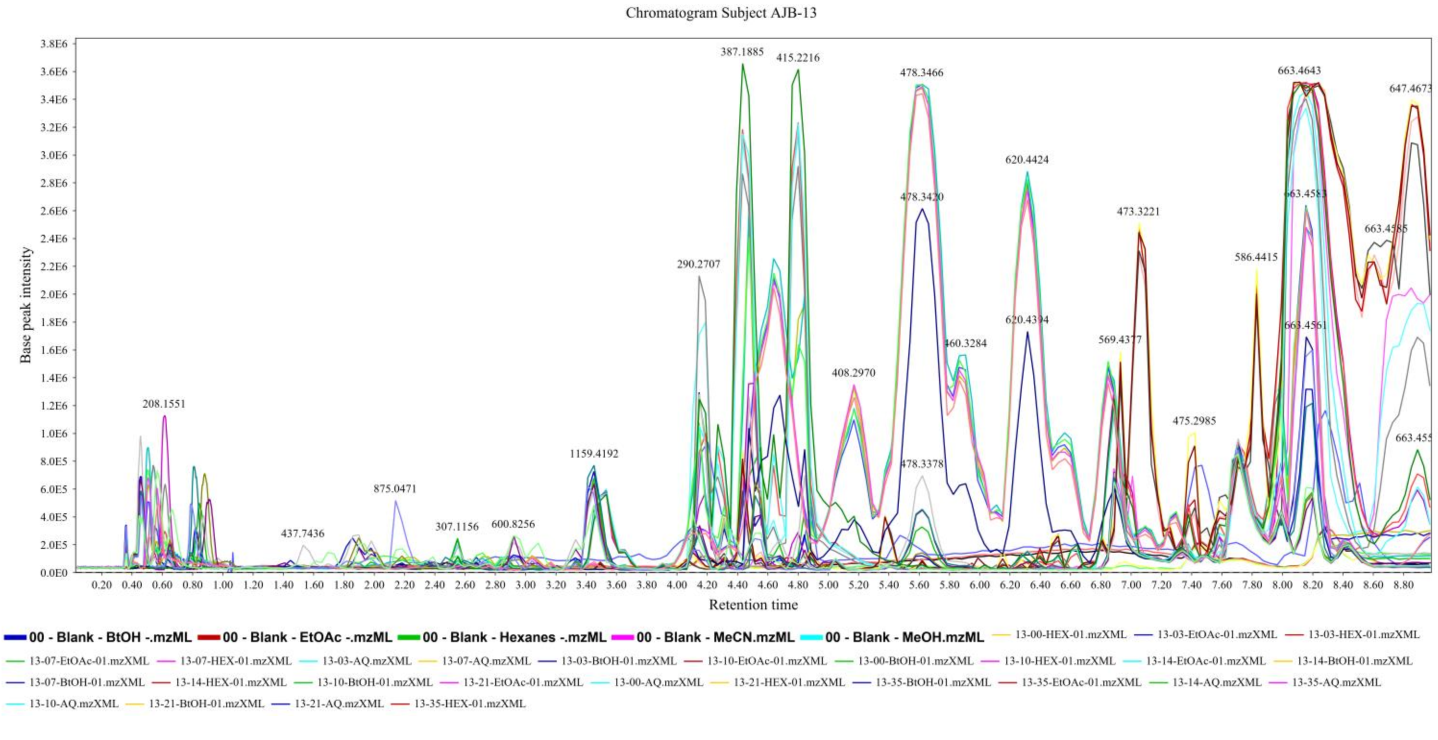


**Supplementary Figure 14. HPLC Chromatogram of Whole Saliva from Subject AJB-13 during Induction and Recovery.**

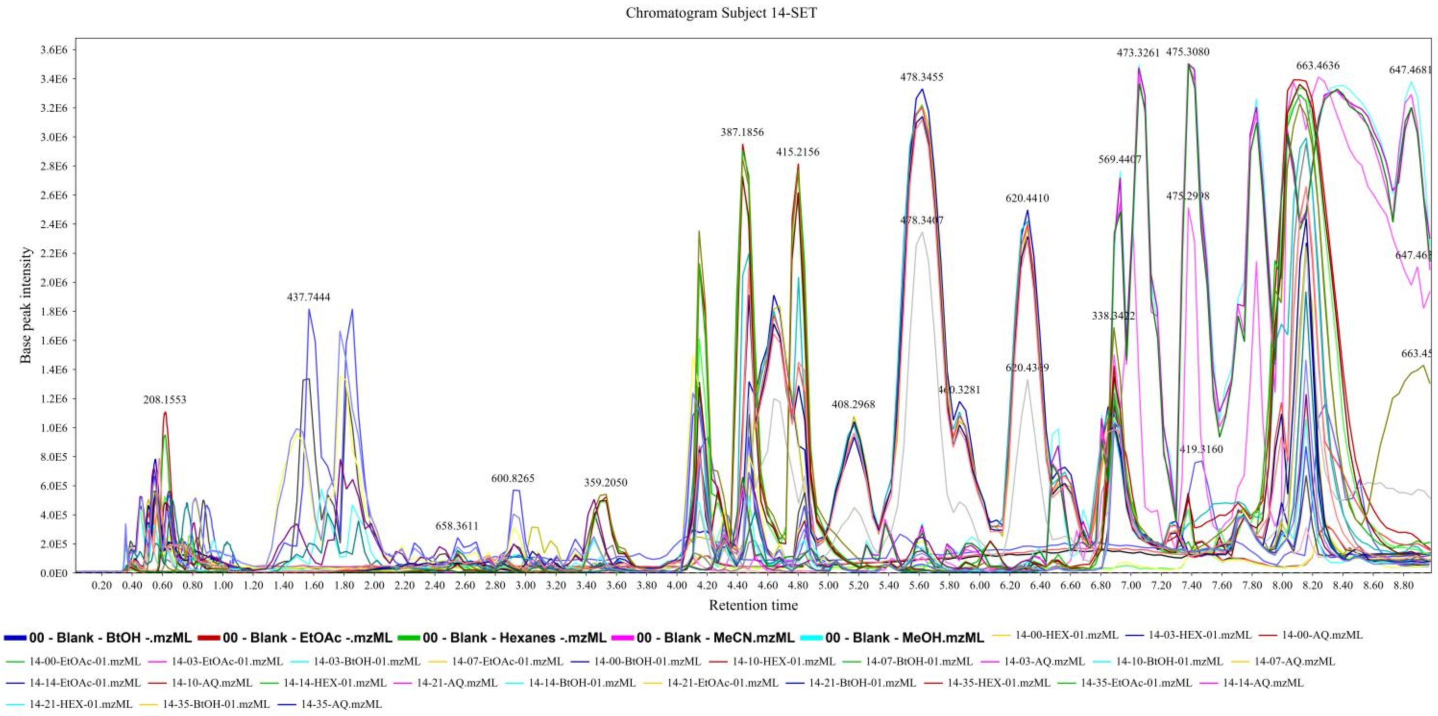


**Supplementary Figure 15. HPLC Chromatogram of Whole Saliva from Subject SET-14 during Induction and Recovery.**

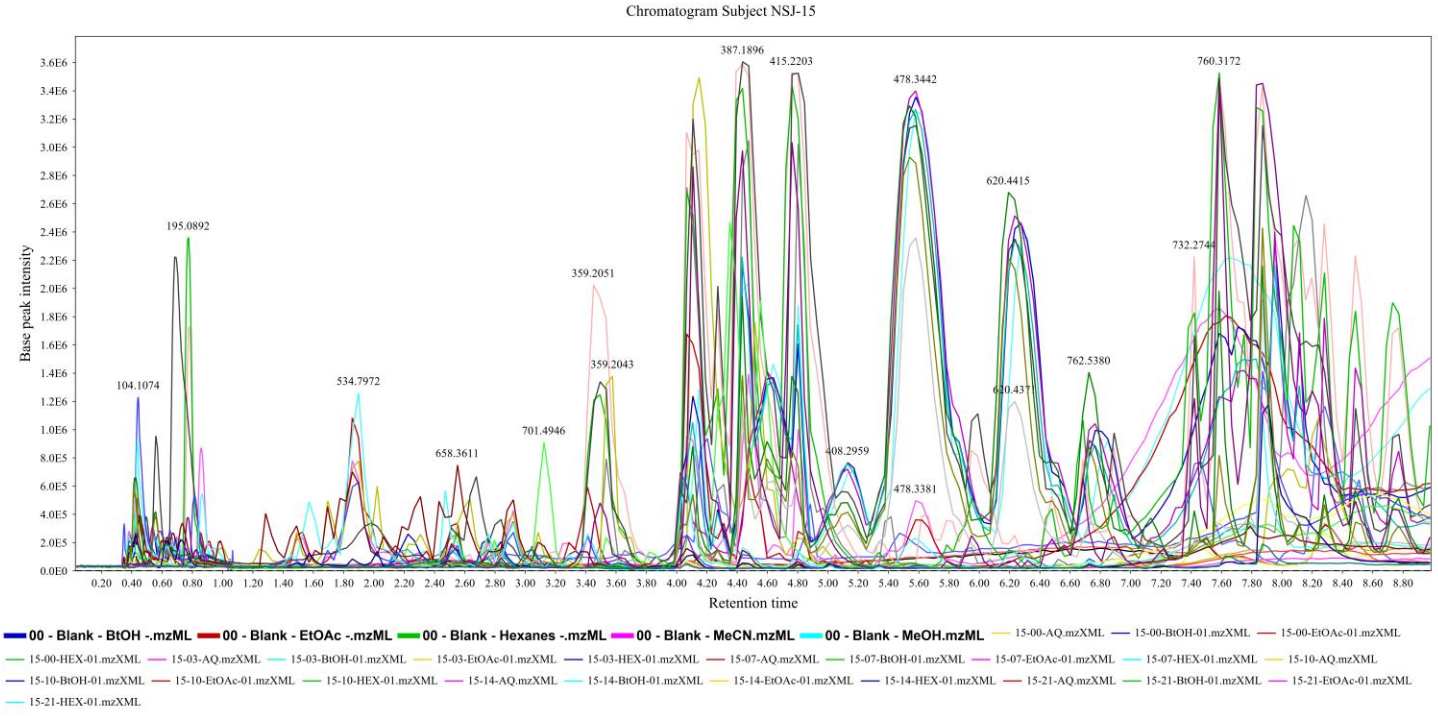


**Supplementary Figure 16. HPLC Chromatogram of Whole Saliva from Subject NSJ-15 during Induction and Recovery.**

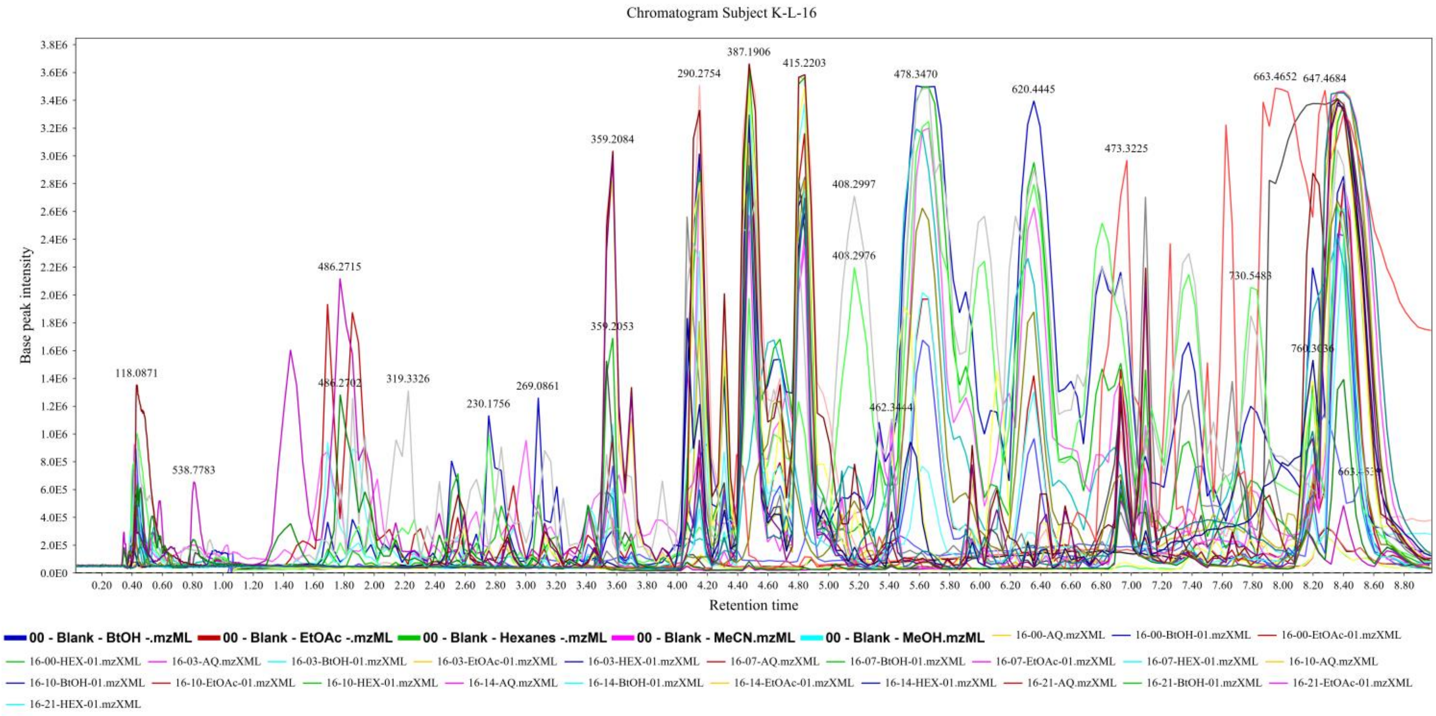


**Supplementary Figure 17. HPLC Chromatogram of Whole Saliva from Subject K-L-16 during Induction and Recovery.**

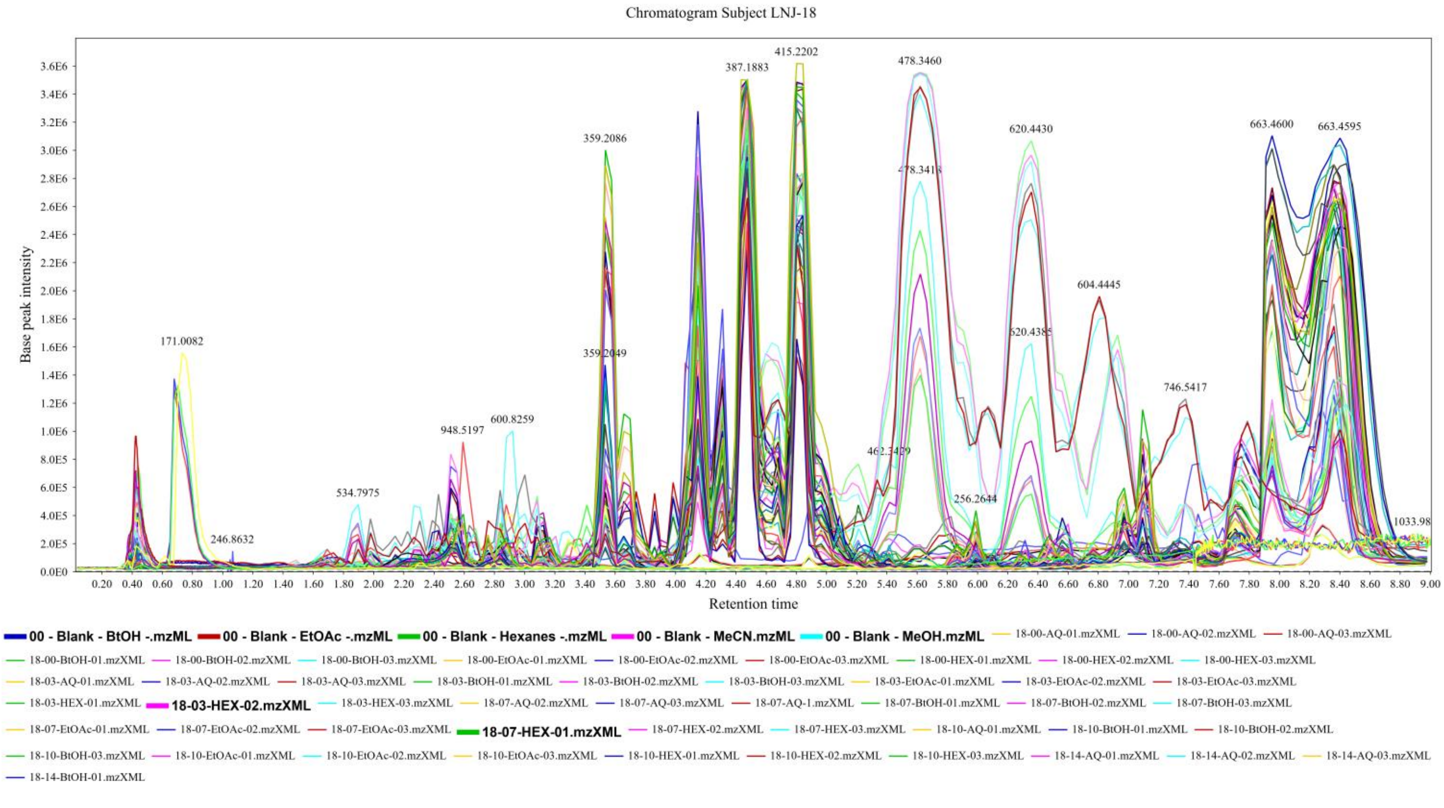


**Supplementary Figure 18. HPLC Chromatogram of Whole Saliva from Subject LNJ-18 during Induction and Recovery.**

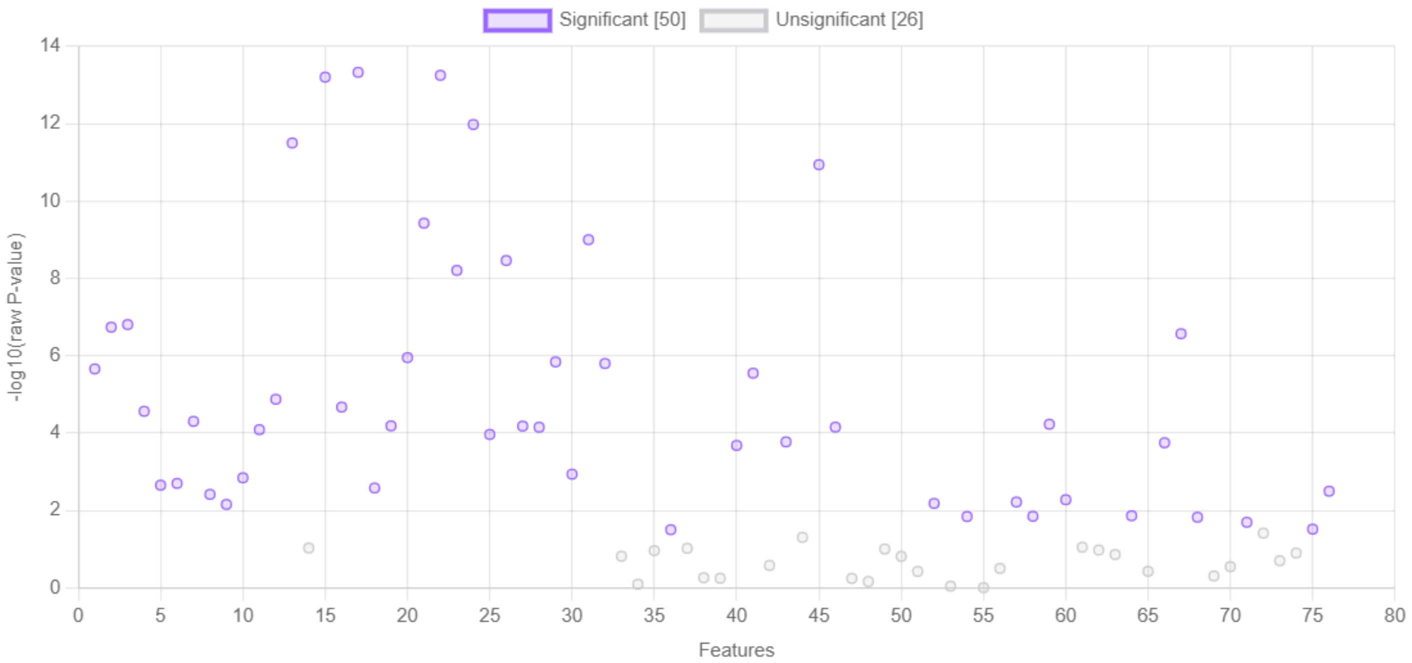


**Supplementary Figure 19.** One-way ANOVA analysis identified 50 statistically significant features modulating throughout the course of gingivitis progression.

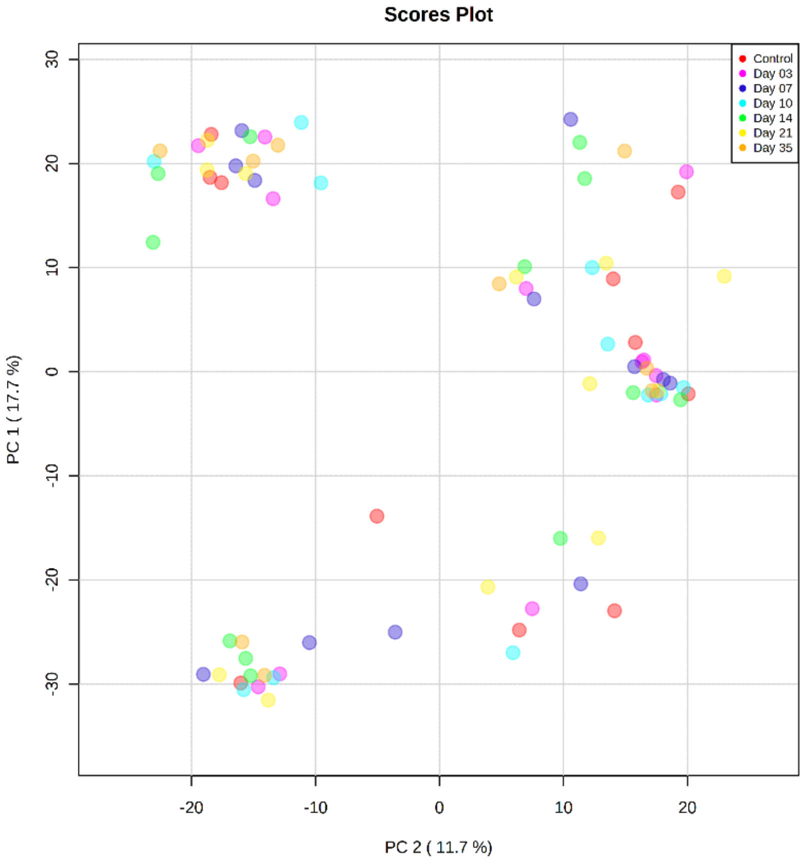


**Supplementary Figure 20. Global Principal Component Analysis of 20 Subject Across Induced Gingivitis.**

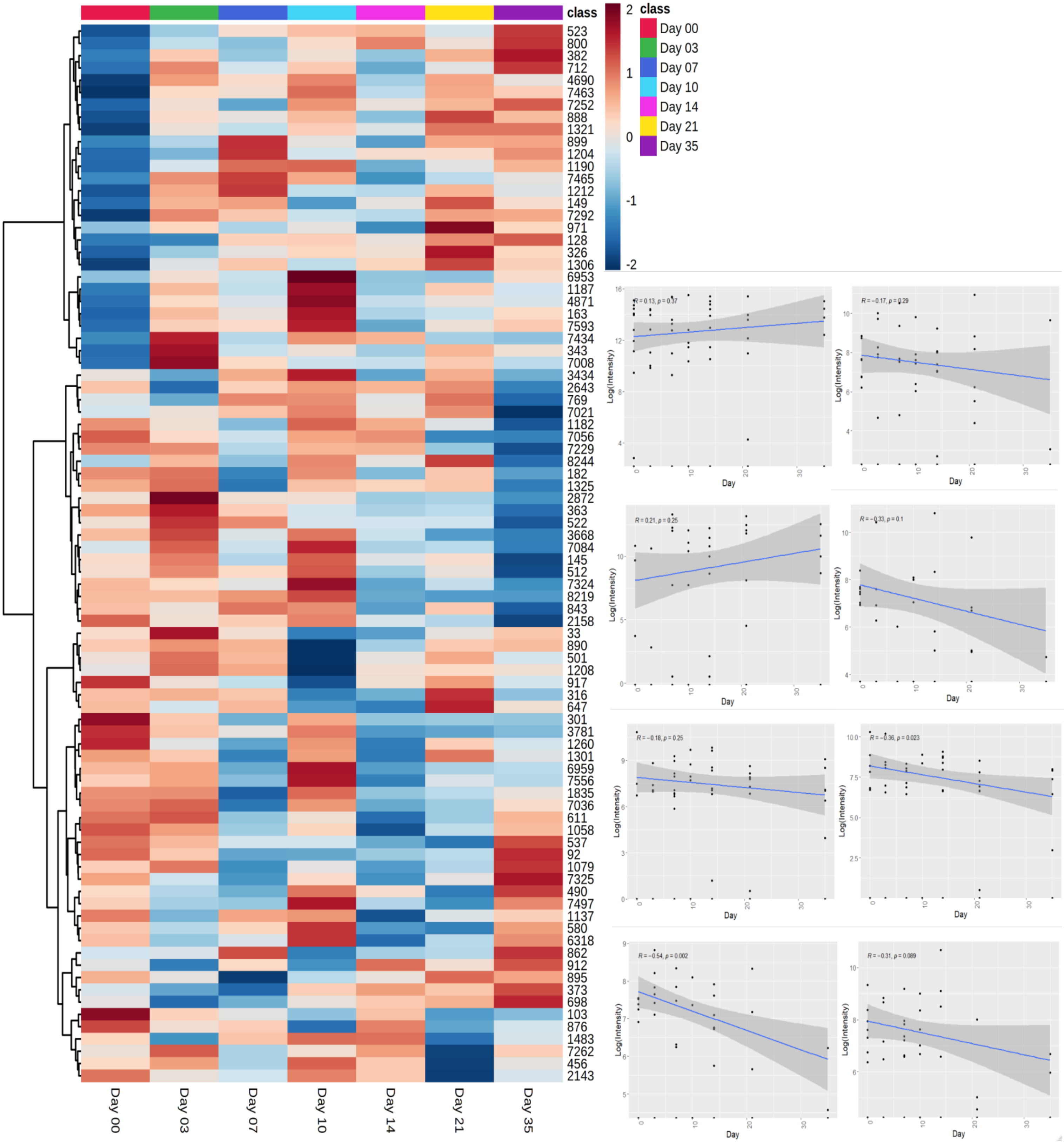


**Supplementary Figure 21. Untargeted Metabolomics for Longitudinal Induced Gingivitis.** Heatmap displays 112 significantly modulating metabolites intensities averaged across individuals. For the 20 subjects within our study, the intensities of eight metabolites were chosen to reveal modulations (upward and downward) due to gingivitis progression.

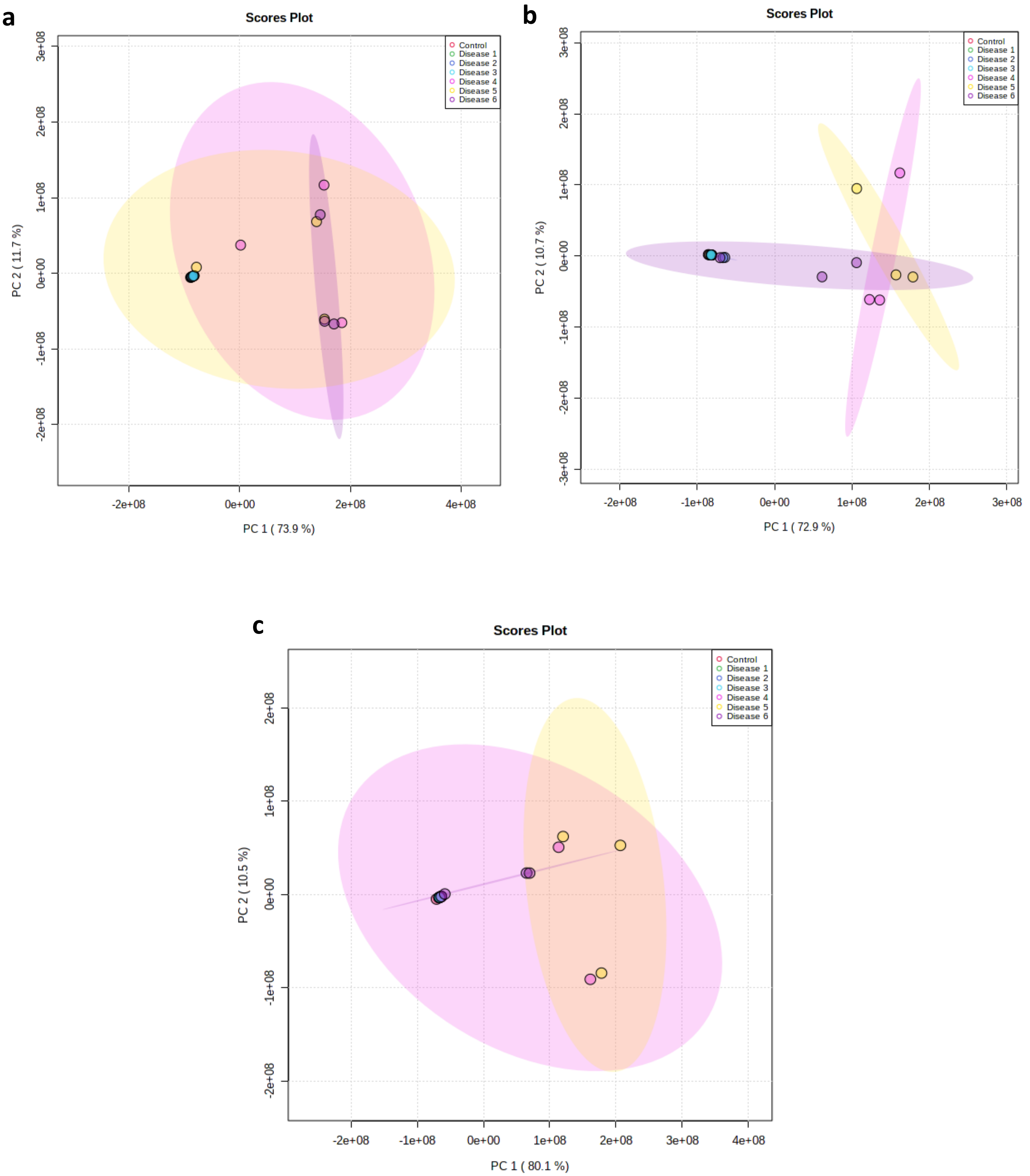


**Supplementary Figure 22A-C. Principal Component Analysis of Subject 3.** Principal component analysis (PCA) distributed individual samples according to the collected time points of EGM (based on 9,651 features). The two PCs explaining the largest part of the variation are shown. **a)** PCA of Hexanes solvent extractions. **b)** PCA of Ethyl Acetate solvent extractions. **c)** PCA of Butanol solvent extractions.

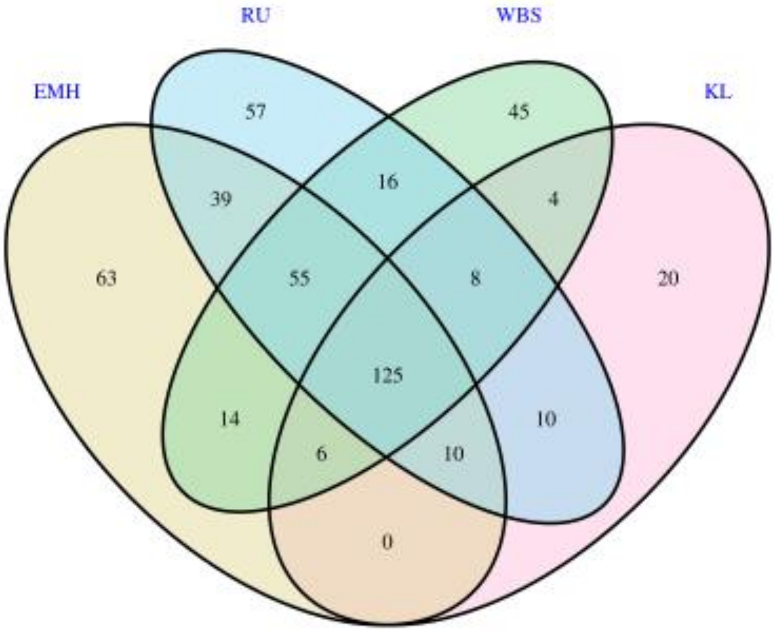


**Supplementary Figure 23. OUT Venn diagram or petal diagram.** The circles of different colors in the Venn diagram represent different samples or groups, and the numbers in the diagram represent the numbers of OTUs unique or common to each sample or group. In the petal diagram, each petal represents a sample or group. The numbers on the petals represent the number of OTUs unique to the sample, and the white circle in the middle represents the number of OTUs shared by all samples and groups.

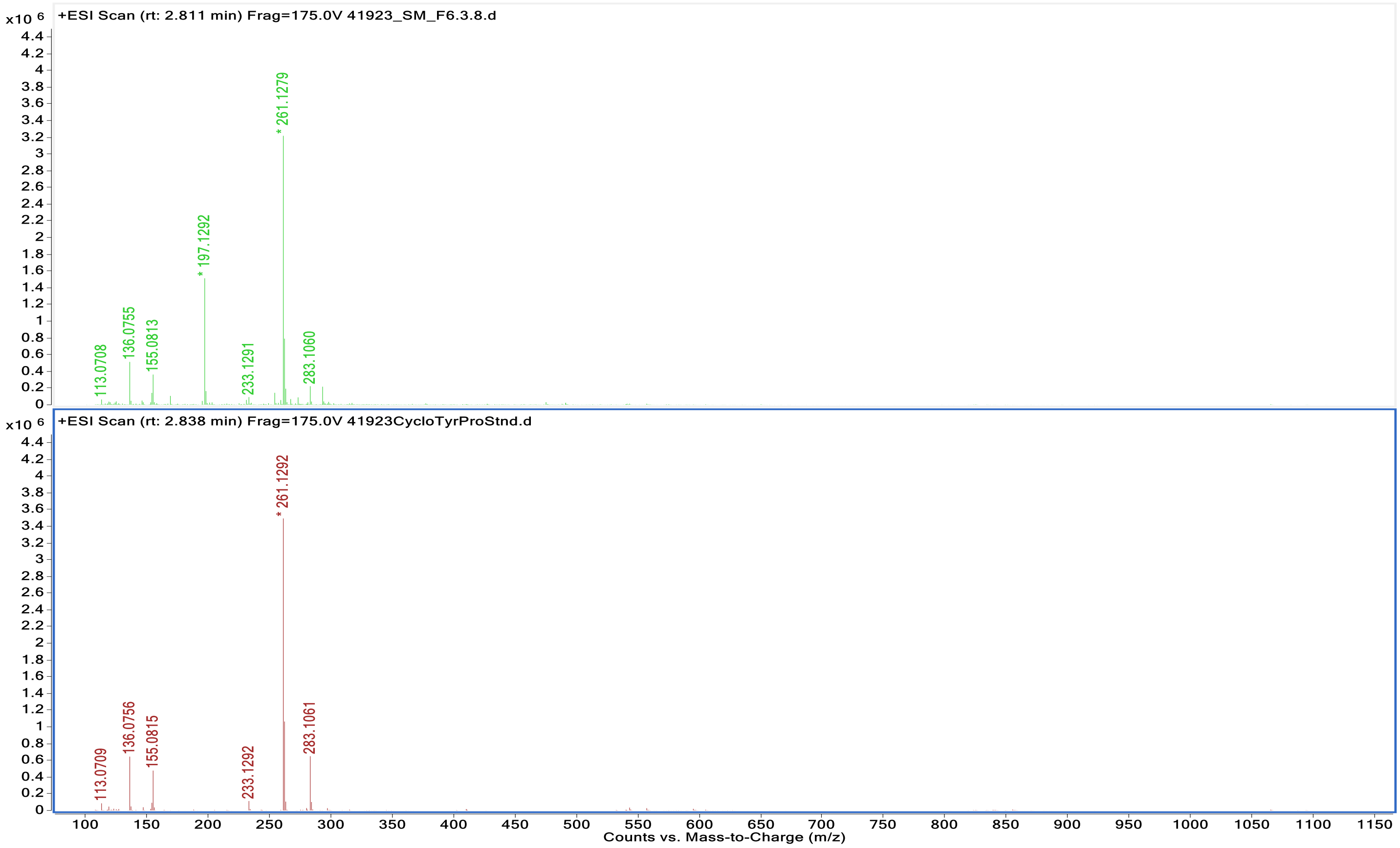


**Supplementary Figure 24**. Mass spectra of DKP, Cyclo(Tyr-Pro), found in S. mitis extract compared to synthetic standard. cavity.

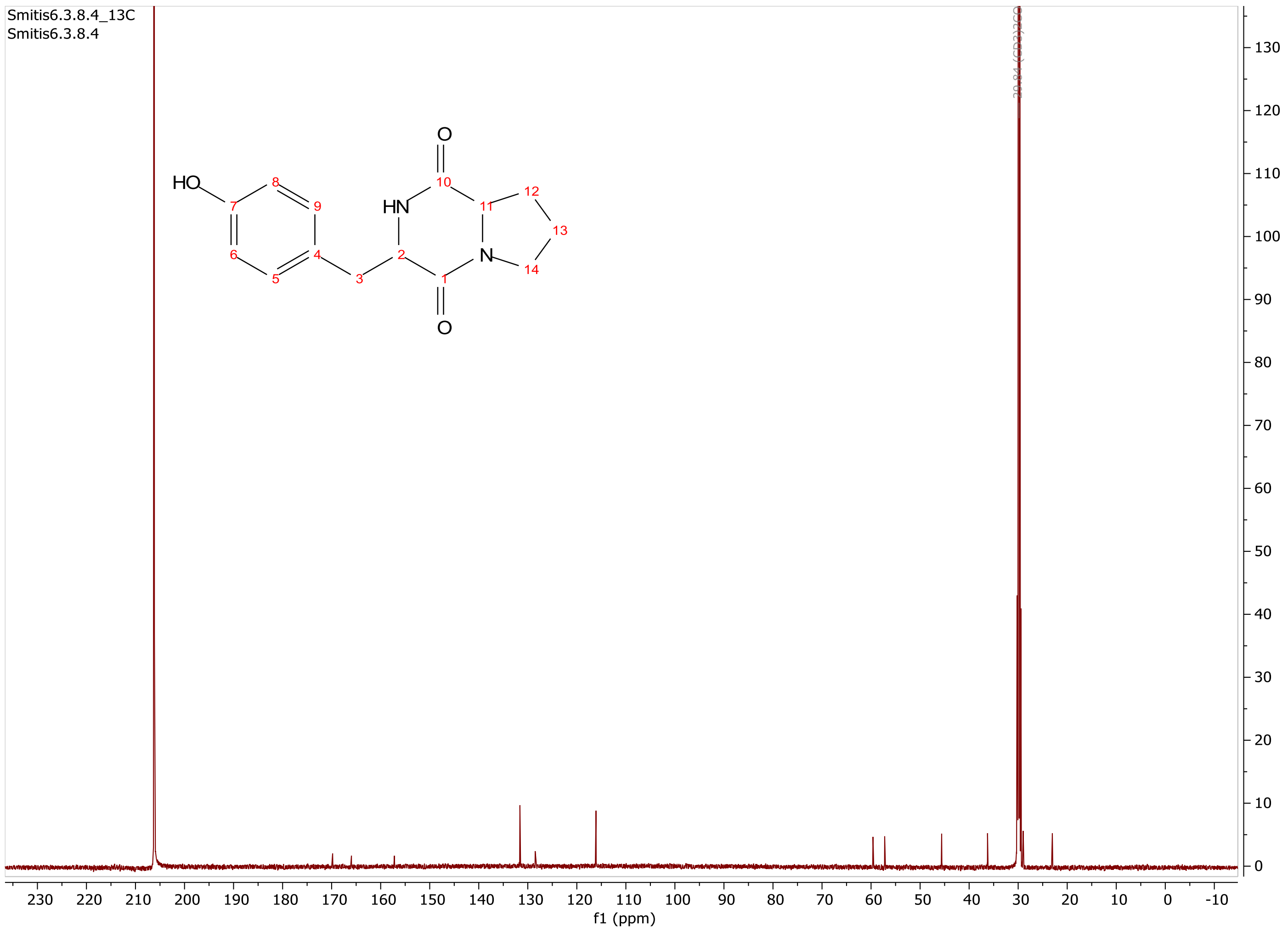


| No. | $\Delta^{13}C$ | $\Delta^{1}H$ (*J* [Hz]) |
|---|---|---|
| 1 | 165.9 | - |
| 2 | 59.6 | 4.29 |
| 3 | 36.2 | 2.91; 3.21 |
| 4 | 128.5 | - |
| 5 | 116.1 | 6.77 |
| 6 | 131.6 | 7.14 |
| 7 | 156.2 | - |
| 8 | 131.6 | 6.77 |
| 9 | 116.1 | 7.14 |
| 10 | 169.8 | - |
| 11 | 57.2 | 4.10 |
| 12 | 28.9 | 2.91 |
| 13 | 23.0 | 1.84 |
| 14 | 45.6 | 3.36; 3.53 |

**Supplementary Figure 25**. $^{13}C$ NMR spectra of DKP, Cyclo(Tyr-Pro) found in S. mitis extract.

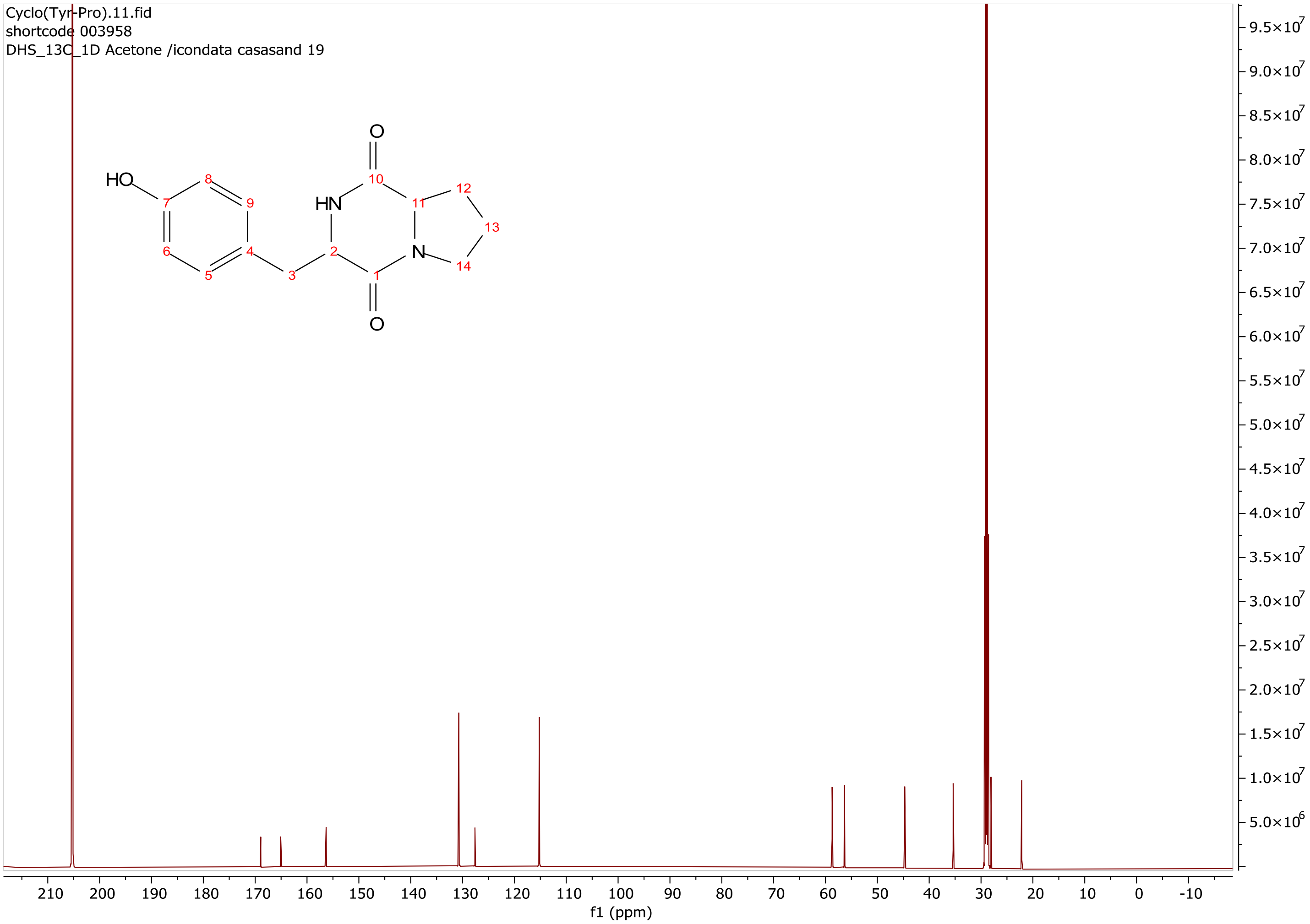


**Supplementary Figure 26**. $^{13}C$ NMR spectra of DKP, Cyclo(Tyr-Pro) synthetic standard.

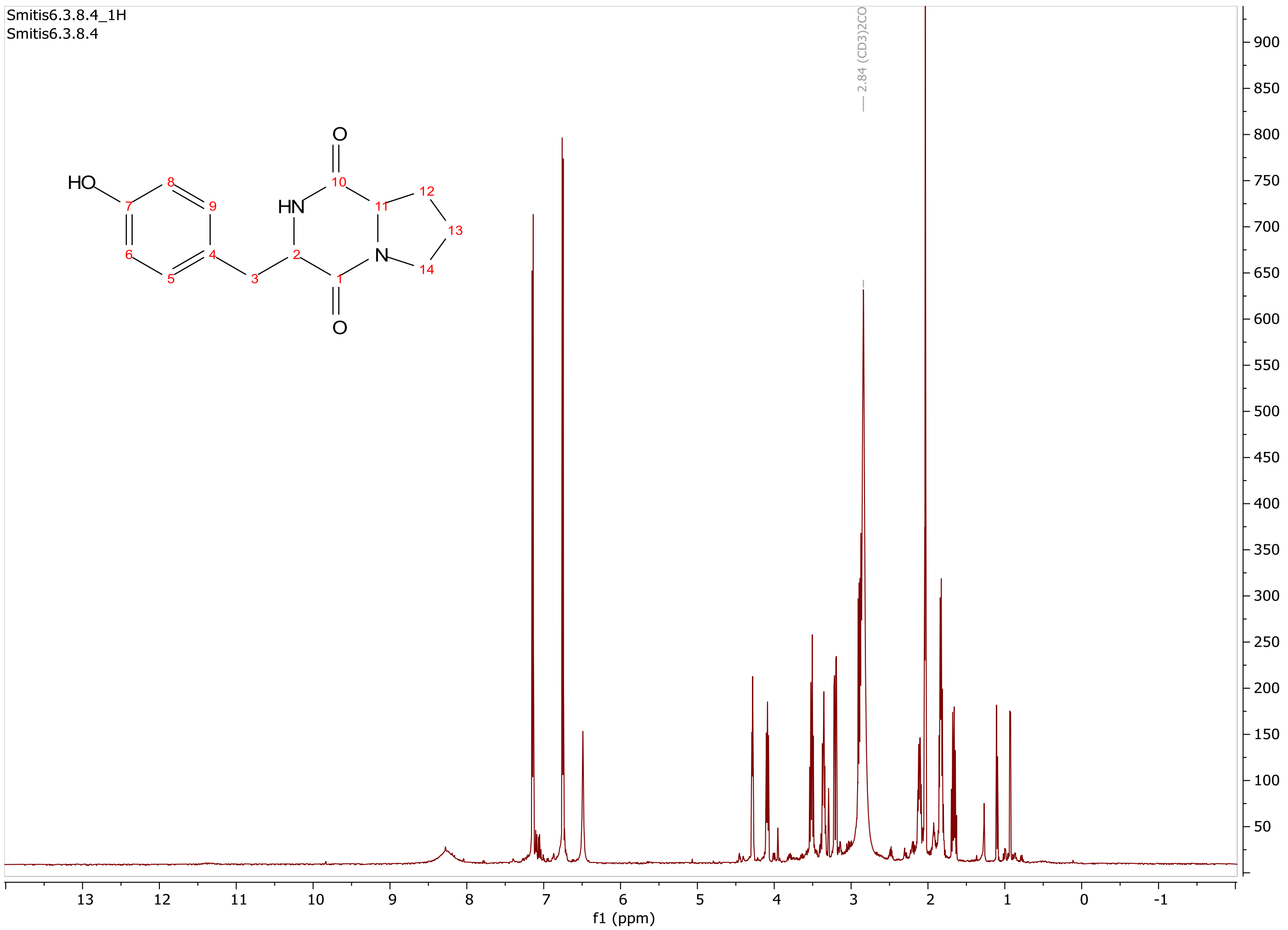


**Supplementary Figure 27**. [1]H NMR spectra of DKP, Cyclo(Tyr-Pro) synthetic standard.

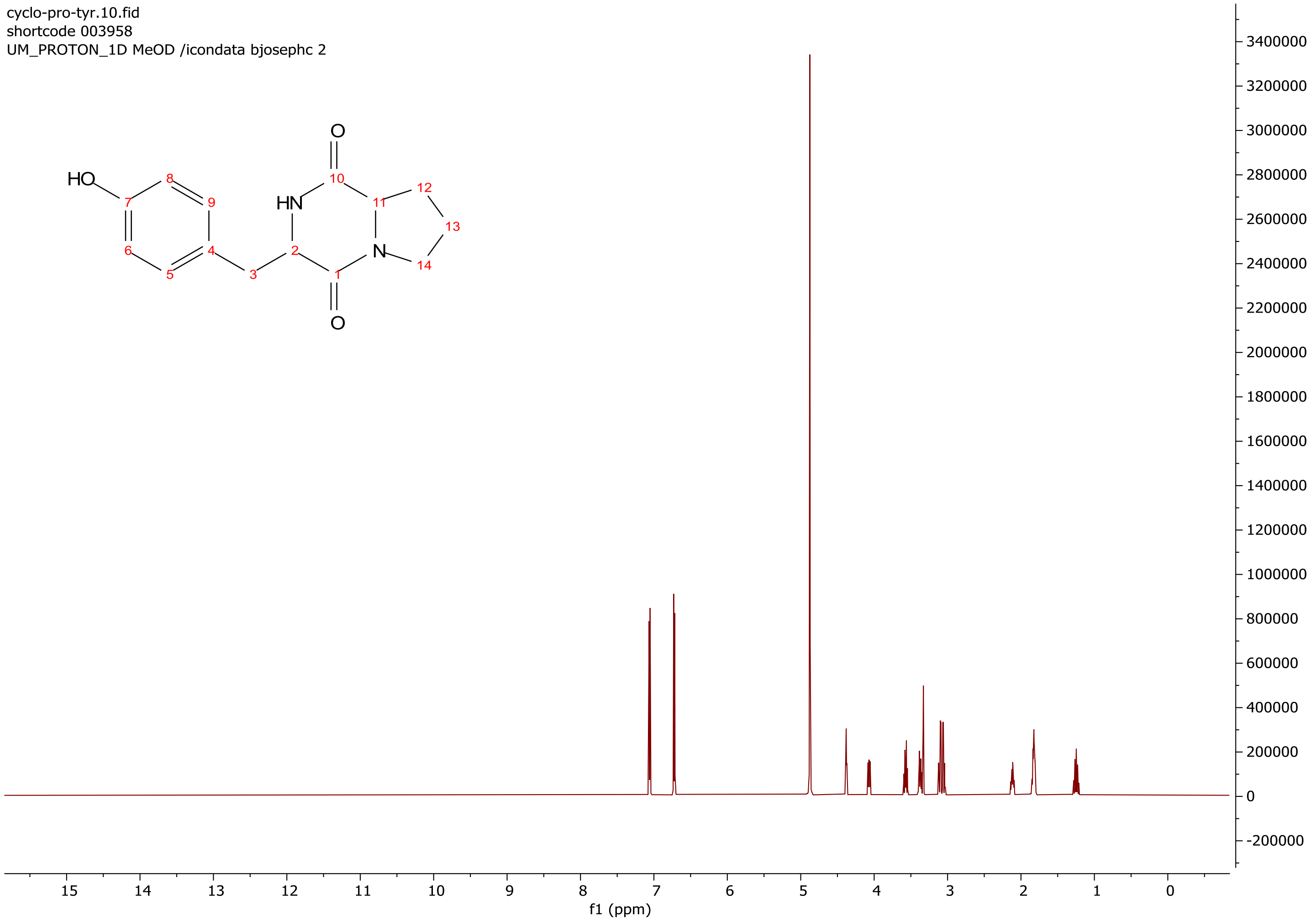


**Supplementary Figure 28**. [1]H NMR spectra of DKP, Cyclo(Tyr-Pro) synthetic standard.
